# A laser capture microdissection-based method for high-sensitivity transcriptomics from archived FFPE tissue slides with single-cell resolution using LCM-FFPEseq

**DOI:** 10.1101/2025.08.13.669096

**Authors:** Elise Callens, Hannes Syryn, Koen Deserranno, Guillaume Cattebeke, Henri Couckuyt, Julie Van De Velde, Danique Berrevoet, Aaron Verplancke, Dieter Deforce, Ward De Spiegelaere, Koen Van De Vijver, Martine Cools, Filip Van Nieuwerburgh

## Abstract

Understanding gene expression within its spatial context is essential for unravelling biological processes. Laser Capture Microdissection (LCM) has emerged as a transformative technology, enabling targeted isolation of individual cells or regions from tissue sections while preserving spatial context. However, its application to formalin-fixed, paraffin-embedded (FFPE) tissues has been limited by RNA degradation, leaving the vast repository of clinical FFPE samples underutilized. To address this, we introduce LCM-FFPEseq, a novel method combining LCM with the advanced Smart-seq3xpress protocol and FFPE-specific adaptations for spatial transcriptomics of FFPE sections. Unlike traditional protocols requiring thousands of cells to generate high-quality libraries, LCM-FFPEseq achieves high sensitivity, reproducibility, and transcript coverage. With as few as 30 FFPE-embedded K562 cells and Sertoli cells, we detected over 14,000 protein-coding genes per sample, with no substantial gains when a higher number of cells were isolated. Even individual LCM-isolated cells yielded an average of 7,353 or 6,490 protein-coding genes per K562 or Sertoli single cell, respectively. To demonstrate its clinical utility, we applied LCM-FFPEseq to archived testicular FFPE samples from transgender females receiving gender-affirming hormone therapy. Transcriptomic profiling of isolated seminiferous tubules revealed tubular hyalinization to be associated with greater upregulation of extracellular matrix remodelling and inflammatory pathways, alongside stronger downregulation of spermatogenesis-associated pathways. These findings suggest that testicular fibrosis and/or tubular hyalinization may contribute to germ cell loss following inappropriate hormonal exposure. By enabling high-resolution transcriptomics in archived FFPE samples, LCM-FFPEseq unlocks new possibilities for investigating rare cell types, spatial heterogeneity, and therapy-induced tissue remodelling in vast FFPE repositories.

## Introduction

Gene expression is regulated both temporally and spatially, making it interesting to study specific cell populations within their native microenvironments to fully understand biological processes and disease mechanisms (Wang, Lin, and Yang 2024). While numerous high-throughput single-cell RNA sequencing (scRNA-seq) methods have been developed, they typically sacrifice spatial information due to tissue dissociation (Li and Wang 2021). Laser capture microdissection (LCM) has emerged as a powerful tool for isolating individual cells or regions of interest from tissue sections with high precision through direct microscopic visualization. This technique overcomes the limitations of whole tissue analysis by enabling the study of distinct or rare cell types specifically, while preserving the histological context and spatial information about the tissue architecture (Datta et al. 2015; Emmert-Buck et al. 1996; Guo et al. 2023; Simone et al. 1998). Moreover, LCM provides advantages over single-cell analysis, which typically requires dissociation of fresh tissue into viable single-cell suspensions, a process that can introduce cellular stress, death, and/or aggregation. This is particularly challenging for certain cell types, such as neurons, which are difficult to isolate intact (Williams et al. 2022).

LCM has been successfully coupled with downstream RNA-seq, enabling precise transcriptomic analysis of specifically selected cells or regions of interest through direct microscopic visualization (Chen et al. 2017; Foley et al. 2019; Nichterwitz et al. 2016). While spatial transcriptomics is a rapidly evolving field, LCM-based RNA-seq remains an established standard due to its contact-free approach, ability to achieve unbiased profiling, and capacity for single-cell resolution (Guo et al. 2023). In contrast, other spatial transcriptomic techniques face drawbacks such as high costs, low resolution, and reduced sensitivity (Guo et al. 2023; Williams et al. 2022). Moreover, these methods are typically optimized for fresh-frozen tissues, making them less suitable for the vast majority of clinically archived samples, preserved as formalin-fixed, paraffin-embedded (FFPE) material (Cheng et al. 2023). Although fresh-frozen tissues are often favoured for transcriptomic applications due to better RNA quality, FFPE tissues are far more abundant, with over millions of FFPE tissue samples collected and archived annually worldwide (Asslaber and Zatloukal 2007; Jin et al. 2024; Vahrenkamp et al. 2019; Waldron et al. 2012). However, FFPE samples present significant challenges for transcriptomic analysis due to crosslinking and RNA degradation. Despite these obstacles, FFPE tissues offer long-term storage at room temperature and superior tissue morphology for identifying specific cell subpopulations (Asslaber and Zatloukal 2007; Evers et al. 2011). To date, formalin remains the standard fixative for tissue preservation in routine clinical practice. Therefore, optimizing LCM-based omics protocols for FFPE samples is highly valuable, as it enables the exploration of spatial heterogeneity and the study of rare cell populations along with their surrounding niche within these unlocked FFPE archives.

Most methods that combine LCM with downstream transcriptomic profiling typically involve a separate RNA extraction step using commercially available RNA isolation kits. These kits not only incur substantial costs but also introduce the risk of sample loss, which is particularly problematic for the already low-input LCM samples. Typically, when LCM is combined with downstream transcriptome analysis using such kits, several hundred to thousands of cells are required as input (Nichterwitz et al. 2016). A significant advancement in this field is LCM-seq (Nichterwitz et al. 2016). This method combines LCM with the Smart-seq2 (SS2) protocol (Picelli et al. 2014), and pioneers in its omission of a dedicated RNA extraction step. By eliminating this step, LCM-seq achieves increased sensitivity, enabling successful processing of samples from as few as five micro-dissected cells, while also substantially reducing overall sample processing time. However, it is important to note that LCM-seq is designed for fresh-frozen tissues and does not allow analysis of FFPE tissue sections (Nichterwitz et al. 2016). Additionally, it does not incorporate certain improvements introduced in later versions of Smart-seq protocols, specifically Smart-seq3 (SS3) (Hagemann-Jensen et al. 2020) and the most recent Smart-seq3xpress (SS3X) (Hagemann-Jensen, Ziegenhain, and Sandberg 2022). These updates include features that could be particularly beneficial for low-input samples derived from LCM. The updated designs for the template-switching oligonucleotide (TSO) and oligo-dT primers in the newer protocols significantly reduce issues related to concatemerization- and strand invasion-artefacts that are particularly problematic in low-input RNA samples such as those obtained via LCM (Hagemann-Jensen et al. 2020, 2022; Picelli et al. 2014). In addition, SS3X incorporates unique molecular identifiers (UMIs), enabling accurate gene expression quantification and mitigation of amplification biases, unlike SS2, which lacks this capability (Picelli et al. 2014). Smart-3SEQ, published in 2019, offers another approach for combining LCM with transcriptomic profiling and is compatible with FFPE samples (Foley et al. 2019). This method integrates the template-switching SMART technique with the 3SEQ protocol for targeting 3’ ends. However, because it builds on SS2 just as LCM-seq, many improvements found in newer protocols are not incorporated, such as modified TSO designs and concentrations to avoid strand invasion, lowering extensively oligo-dT primer concentrations to reduce PCR side product formation, the use of PEG-8000 instead of betaine to boost performance, and the use of GTP within the RT mix to increase sensitivity. These shortcomings are particularly significant when working with low-input RNA, where preventing PCR side products is crucial to minimize adapter dimer formation and improve yield. Consequently, LCM-FFPE samples processed with Smart-3SEQ have shown very low (exon) mapping rates (Foley et al. 2019; Hagemann-Jensen et al. 2020, 2022; Picelli et al. 2014).

To address these challenges, we introduce LCM-FFPE-seq, a method that combines LCM of FFPE tissues with an adapted version of the recent SS3X RNA-seq technology. This approach integrates the enhanced sensitivity of SS3X with FFPE-specific adaptations from the Smart-3SEQ protocol. By omitting a separate RNA extraction step, LCM-FFPEseq streamlines sample processing and improves cDNA yield from low-input material. In addition, the SS3X protocol includes UMIs for error-correction and incorporates a biotin-modified TSO to prevent concatemerization, ensuring higher accuracy and alignment rates in LCM-FFPE samples. To validate the LCM-FFPEseq method and evaluate its sensitivity and robustness, we performed a controlled dilution series of microdissected FFPE-embedded K562 cells, ranging from single cell to 500-cell inputs, allowing us to define the suited working range. To demonstrate the clinical utility, we applied LCM-FFPE-seq to archived, FFPE-preserved testicular tissue from transgender females to investigate the effects of gender-affirming hormone therapy (GAHT) at the tissue level. This setting uniquely benefits from LCM-FFPE-seq, as it allows for the selective capture of seminiferous tubules and enables the use of archived samples that are not commonly available.

Adolescents registered male at birth and who pursue female transition receive prolonged treatment with GnRH analogues or cyproterone acetate (CPA) for puberty suppression, followed by the addition of estrogen treatment, starting from the age of 16 years onwards (Coleman et al. 2022). In this study, the combination of CPA and estrogen treatment will be referred to as GAHT. Using this treatment, testes are exposed to exogenous hormones for many years, prior to gonadectomy, which is usually performed after the age of 18. The impact of these medications on gonadal histology and function remains poorly understood, and is important as some trans females opt to maintain their gonads, as they may wish to produce mature sperm for reproductive purposes. Histological changes such as tubular hyalinization with basement membrane thickening, and absence of gametes have been observed in some trans female individuals (Cornejo et al. 2022; de Nie et al. 2022). Specific capture of Sertoli cells, being the germ cell-supporting cells, can shed further light on the underlying mechanisms of these changes. The ability to use these archived samples enables the investigation of therapy-induced tissue remodelling and potential damage, as will be illustrated below.

In summary, LCM-FFPEseq opens new possibilities for transcriptomic profiling of archived FFPE samples, allowing for a more precise exploration of gene regulatory networks and signalling pathways, even in rare cell populations and spatially distinct tissue regions.

## Results

### LCM-FFPEseq workflow

Our LCM-FFPEseq assay couples LCM with the design of the recent SS3X protocol, however with FFPE-specific Smart-3SEQ optimizations, as displayed in Figure 1 and the detailed schematic overview of the library preparation in Supplementary Figure 1. FFPE tissue slides are sectioned at 7 µm, mounted on polyethylene naphthalate (PEN) membrane glass slides, and stained using the Applied Biosystems™ Histogene™ (Arcturus) quick staining, specifically tailored for LCM coupled to RNA-seq application. The staining provides good contrast by differential staining of nuclei (purple) and cytoplasm (light pink) and preserves RNA integrity due to its brief exposure time of only 30 seconds. Similar to LCM-seq, we combine the Histogene™ staining with off-the-shelf ethanol to eliminate the need to purchase expensive commercial staining kits. However, FFPE tissue sections require an initial xylene step for paraffin removal, followed by a decreasing ethanol series for xylene removal and tissue rehydration. After Histogene™ staining, the tissues undergo an increasing ethanol series for tissue dehydration. The common final xylene step is omitted and air-drying only was performed to assure the adhesion of the tissue to the PEN membrane glass slide (PALM Protocols-DNA Handling n.d.). Alternative (antibody) staining methods can be employed to target specific cell types or biomarkers; however, quality control is essential, as certain staining protocols may have a more pronounced detrimental impact on RNA quality than others (Clément-Ziza et al. 2008; Wang et al. 2006).

**Figure 1.**
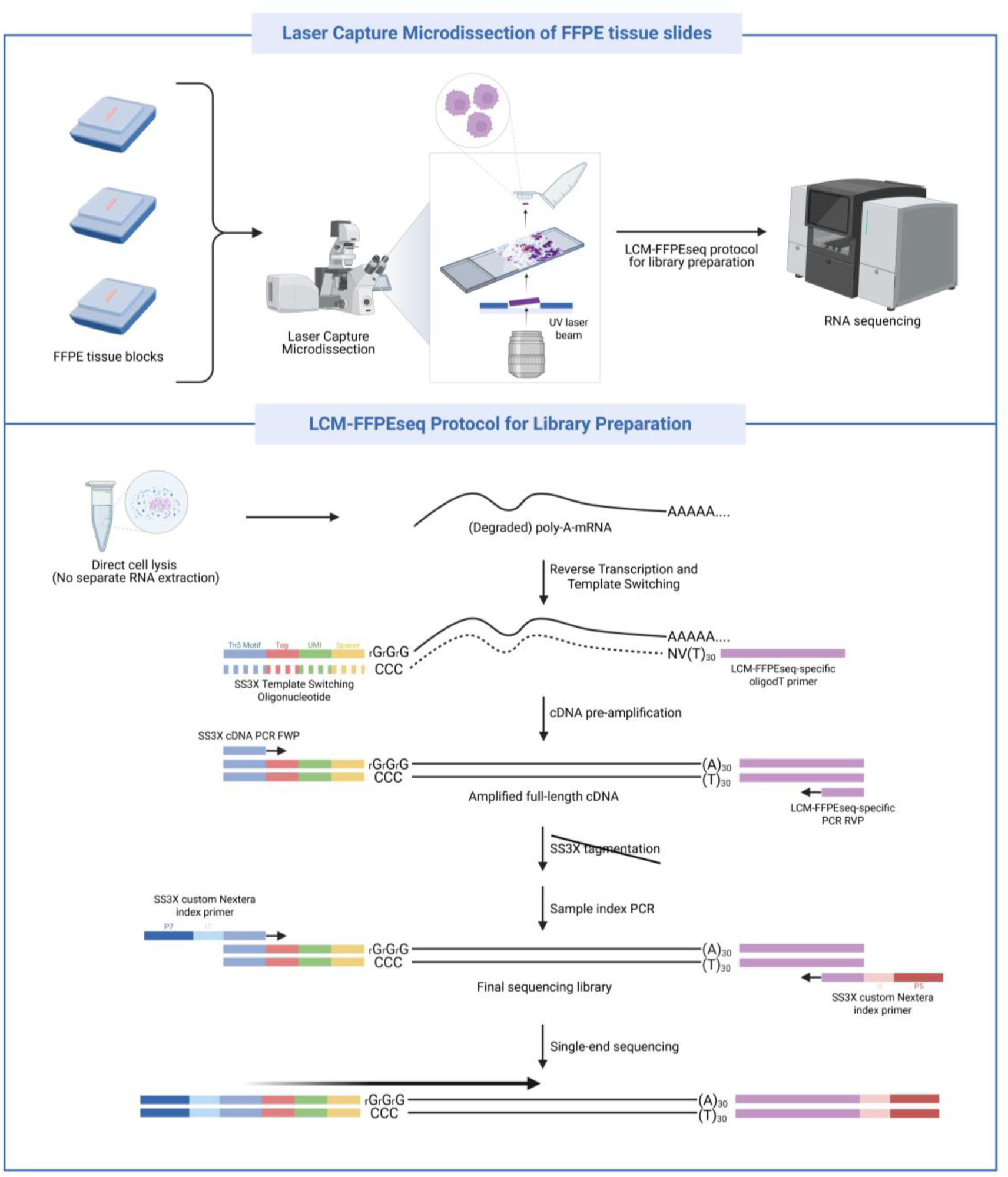
Overview of the LCM-FFPEseq workflow. FFPE tissue blocks were sectioned at 7 µm thickness, mounted on PEN membrane glass slides and stained using Histogene™. Cells or regions of interest were visualized using the PALM MicroBeam LCM System (ZEISS), then precisely microdissected and catapulted into adhesive cap PCR tubes. Library preparation was performed using the LCM-FFPEseq protocol, which is based on the SS3X protocol and incorporates FFPE-specific adaptations, including a direct lysis step with proteinase K that eliminates the need for a separate RNA extraction. Following lysis, reverse transcription, cDNA amplification, and sample indexing were carried out. Final sequencing was performed in single-end mode.

A separate RNA extraction is omitted, and direct lysis is performed on the LCM-isolated cells, rendering a time-efficient and highly sensitive protocol. Following microdissection, the cells are dry-shot into adhesive cap PCR tubes, allowing the lysis buffer to be added directly when initiating library preparation. However, since the used SS3X lysis buffer is not specifically suited for FFPE tissue, FFPE-specific adaptations are implemented. Therefore, micro-dissected FFPE tissues are incubated for 50 minutes at 52°C in the presence of proteinase K-lysis buffer to lyse the tissue, followed by a heating step of 10 minutes at 72°C to inactivate the enzyme. In addition, a proteinase K inhibitor is included in the reverse transcription (RT) mix to further promote proteinase K inhibition after lysis, as this complementary approach results in a large improvement in library yield. It should be noted that in contrast to the miniaturized SS3X protocol designed for single FACS-sorted cells, our method is not equally miniaturized. Volumes are scaled up to allow for manual pipetting and compatibility with LCM-isolated material, typically processed in 0.2 mL adhesive cap PCR tubes.

Following lysis, RT is performed using an LCM-FFPEseq-specific oligo-dT primer. Due to the inherently degraded nature of FFPE material, tagmentation as performed in SS3X is unsuitable, as it would generate an excessive number of reads that are too short for reliable mapping. Therefore, the oligo-dT primer is designed to introduce binding sites for both PCR pre-amplification and final sequencing adaptors. After first-strand cDNA is pre-amplified via PCR, a second amplification step is conducted to incorporate the sample indices and sequencing adaptors. Importantly, since no fragmentation step is performed, the poly-A tail remains intact. This grounds the choice for single-end sequencing of the final sequencing library. Generated LCM-FFPEseq sequencing data is downstream compatible with the zUMIs pipeline, which is typically used for processing standard SS3X data.

### Validation of LCM-FFPEseq method

To validate the developed LCM-FFPEseq method, we utilized FFPE-embedded K562 cells to assess the effectiveness of the direct lysis, RT, and PCR, as well as to evaluate the method’s sensitivity and reproducibility in a controlled, homogeneous setting. Given that LCM procedures can benefit from minimized input requirements, both to save time and to accommodate cases where tissue or cells of interest are scarce, we explored the minimum cellular input needed to generate high-quality RNA-seq libraries. Working with a separate RNA extraction kit would typically require thousands of LCM-isolated cells, however, our experimental design involved isolating 500 cells across multiple iterations with LCM and progressively scaling down to 300, 100, 50, 30, 10, 5, and ultimately a single cell (Supplementary Figure 2). For the non-single-cell samples, cells were isolated in group rather than individually. In addition, two negative controls were included. To evaluate cDNA yield and quality following direct lysis and RT, all samples were subjected to 21 cycles of PCR pre-amplification. The resulting cDNA profiles were observed using the Agilent Fragment Analyzer (Supplementary Figure 3), and cDNA yield was determined by Qubit Fluorometric Quantification. Table 1 summarizes the number of replicates, the total area of collected cells (µm²), and the total cDNA yield (ng) for each group.

**Table 1.** Specifications of dilution series of LCM-isolated FFPE-embedded K562 samples and corresponding cDNA libraries. Values are mean ± SD. NA, not applicable. *Extra (blank) margin was taken to avoid damage of laser to single cell.

| **<u>Number of LCM-isolated cells</u>** | **<u>Replicates</u>** | **<u>Cells collected</u>** | **<u>Total Area (μm<sup>2</sup>)</u>** | **<u>cDNA yield (ng)</u>** |
| --- | --- | --- | --- | --- |
| 0 cells | 2 | NA | NA | 11.6 ± 0.3 |
| 1 cell | 3 | 1 ± 0 | 2,115 ± 253* | 11.4 ± 2.3 |
| 5 cells | 3 | 5 ± 0 | 1,823 ± 597 | 11.1 ± 4.6 |
| 10 cells | 3 | 10 ± 0 | 4,307 ± 369 | 9.4 ± 1.1 |
| 30 cells | 3 | 30.7 ± 0.6 | 8,810 ± 1,898 | 25.4 ± 3.3 |
| 50 cells | 3 | 50.7 ± 0.6 | 13,922 ± 976 | 24.0 ± 6.3 |
| 100 cells | 3 | 102 ± 1.7 | 30,397 ± 2,510 | 35.3 ± 9.3 |
| 300 cells | 3 | 300.9 ± 3.8 | 79,957 ± 3,916 | 46.7 ± 9.3 |
| 500 cells | 3 | 498.3 ± 0.6 | 116,369 ± 1,430 | 45.8 ± 16.7 |

Sequencing results revealed a success rate of 67% for single-cell samples, and 100% for samples with five or more cells. Detected gene counts are illustrated in Figure 2A, and a detailed overview of RNA-seq metrics is provided in Supplementary Table 1. The LCM-FFPEseq protocol exhibited high sensitivity and reproducibility for inputs of 30 up to 500 cells, detecting an average of 16,246 protein-coding genes per sample. The highest gene detection was observed at the 300-cell input, with an average of 17,098 protein-coding genes identified. The total UMI count ranged from 50,019 for successful single cells to on average 2,814,840 for 500 cell-samples. While the protocol remained successful for inputs below 30 cells, reproducibility and gene counts were reduced. However, successful single-cell samples still achieved an average gene count of 7,353 genes. Total mapping ratios improved with cell input, ranging from 25.7% for successful single-cell samples to 76.1% for samples with 500 cells. Exon mapping ratios followed a similar trend, increasing from 8.7% for successful single cells to 35.9% for 500 cells. Among all read types, exon mapping reads were the most abundant, followed by unmapped, intergenic, intronic, and ambiguity reads (Figure 2B). To further explore the types of transcripts detected across the dilution series, we analysed gene biotypes according to Ensembl annotations (Figure 2C). Protein-coding genes were the most frequently detected, reflecting the poly(A)-specific enrichment using oligo-dT priming. However, a substantial fraction of long non-coding RNAs (lncRNAs) was also identified, likely poly(A)-containing lncRNAs, as well as some pseudogenes. Gene body coverage analysis revealed a pronounced 3’ bias (Figure 2D), consistent with the protocol’s reliance on poly(A)-tail priming and the absence of a tagmentation step. Library complexity was evaluated across the dilution series using saturation curves plotting the detected unique UMI counts against the total number of reads per replicate (Supplementary Figure 4). While single-cell and low-input samples (1– 10 cells) showed more variability between replicates and did not reach full saturation, higher-input samples (≥30 cells) exhibited consistent complexity profiles and reached saturation across the replicates. Principal Component Analysis (PCA) revealed three clusters (Figure 2E). The first cluster, located in the upper left quadrant, includes the blank samples, specifically the negative controls and one failed single-cell capture. The second cluster, positioned in the lower central area, comprises the successful low-input samples (1–10 cells). The third and most compact cluster, centrally located, contains all mid- to high-input samples (≥30 cells). Gene expression correlation was high within the third cluster, whereas low-input samples, despite originating from identical cells, exhibited greater variability across replicates. The sample-to-sample correlation heatmap (Figure 2F) further supports these observations, showing strong correlation among mid- and high-input samples, with a gradual decrease in similarity as input size decreases.

**Figure 2.**
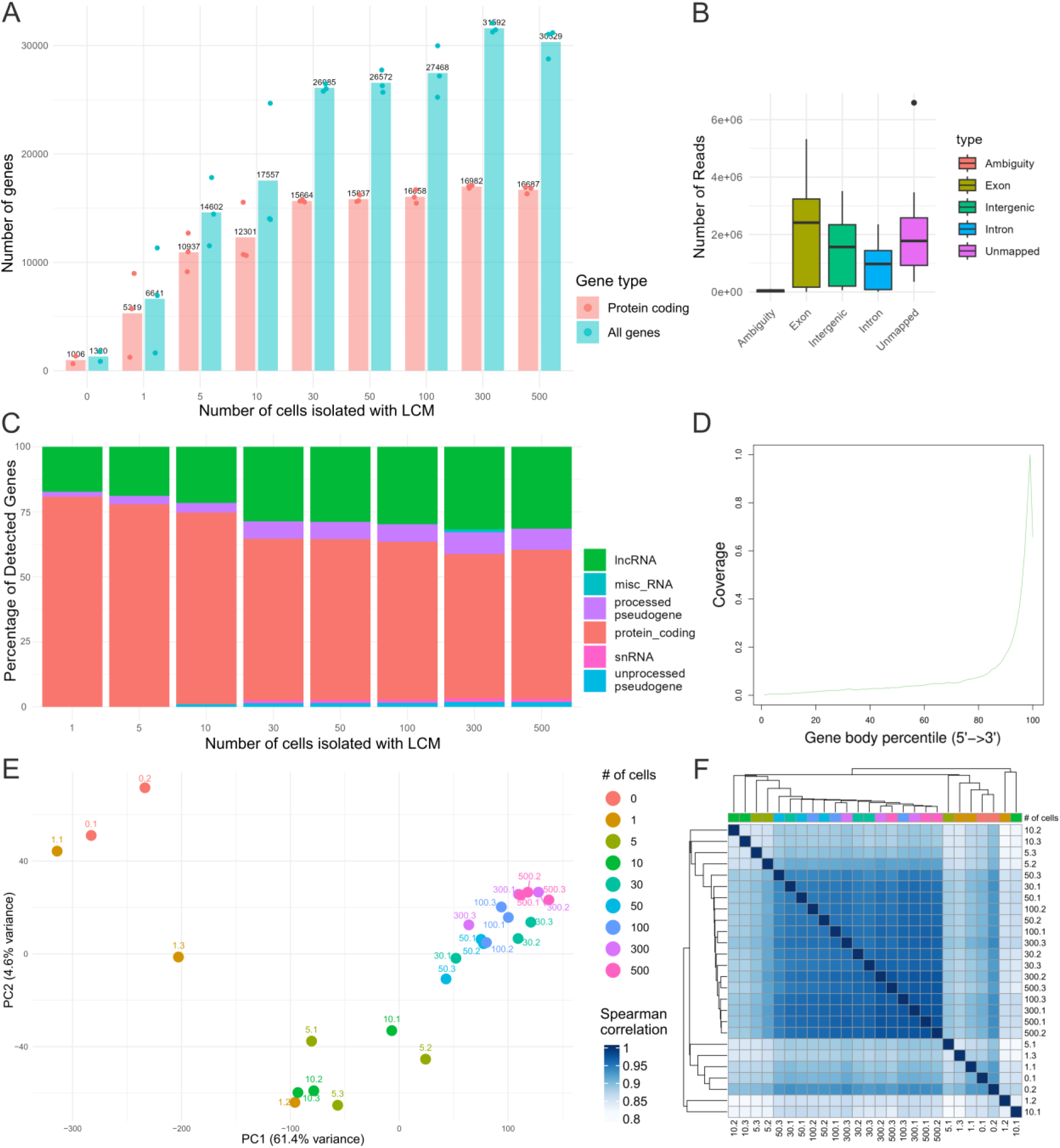
LCM-FFPEseq dilution series results of LCM-isolated FFPE-embedded K562 cells. (A) Average number of detected genes across varying input levels, shown separately for protein-coding genes and all detected genes. Individual replicates are shown as dots, while bars represent the mean gene count per condition. (B) Average read distribution across different genomic feature categories: ambiguity, exon, intergenic, intron and unmapped reads. (C) Proportional average gene biotype distribution per group. (D) Gene body coverage analysis across the 5’ to 3’ regions. (E) Unsupervised two-dimensional PCA plot for all samples of the dilution series. Sample IDs are indicated as *X.Y*, where *X* represents the number of cells isolated with LCM and *Y* indicates the replicate number. (F) Sample-to-sample correlation heatmap of expression profiles across all samples of the dilution series, based on Euclidean distance and hierarchical clustering. Sample IDs are indicated as *X.Y*, where *X* represents the number of cells isolated with LCM and *Y* indicates the replicate number.

### LCM-FFPEseq on archived human testicular FFPE tissue sections

#### Clinical and Histopathological Data

LCM-FFPEseq was applied on testicular tissue to investigate potential mechanisms underlying alterations associated with GAHT. A summary of the sample characteristics can be found in Table 2. Six testicular FFPE samples were selected from transgender females (individuals registered male at birth) who underwent gonadectomy following suppression of endogenous testosterone production via CPA, followed by estrogen administration during puberty to induce bodily feminisation. The mean age at initiation of CPA administration was 16.4 years (range: 14.9-18.3 years), and for estrogen, 17.2 years (range: 16.0-18.6 years). The average duration of CPA treatment was 2.6 years (range: 1-4.3 years), and of estrogen treatment, 1.8 years (range: 0.8-3.2), until gonadectomy was performed at a mean age of 19 years (range: 18.1-19.8 years). Histological analysis revealed marked tubular hyalinization in three samples, while the remaining three showed no such tubular hyalinization (Figure 4A). All testes displayed mature Sertoli cells and the presence of spermatogonia, with a Johnsen score (i.e. a measure for the maximal spermatogenic stage reached in a certain sample, with a maximum of 10) of 3. As controls, testicular FFPE tissue was included from two cisgender males aged 17 and 23 years with histologically confirmed normal spermatogenesis. The FFPE samples were archived between 6 months and 4 years. To validate LCM-FFPEseq in a biological context, control tissue (Sample 1) was used for a dilution series of Sertoli cells and to compare two different cell types (Sertoli and Leydig cells) at the single-cell level. Finally, transcriptional differences were assessed between GAHT-exposed testicular tissues and those from untreated controls using larger LCM-isolated areas comprising individual seminiferous tubules (± 200,000 µm²).

**Table 2.** Characteristics of included testicular samples. CPA: cyproterone acetate, E2: estrogen, +: yes, -: no, NA: not applicable.

| <u>Sample</u> | <u>Age at start</u><br><u>CPA (years)</u> | <u>Age at start</u><br><u>estrogen (years)</u> | <u>Age gonadectomy</u><br><u>(years)</u> | <u>Duration of CPA</u><br><u>exposure (years)</u> | <u>Duration of E2</u><br><u>exposure (years)</u> | <u>Tubular</u><br><u>hyalinization</u> |
| --- | --- | --- | --- | --- | --- | --- |
| 1 | NA | NA | NA | NA | NA | - |
| 2 | NA | NA | NA | NA | NA | - |
| 3 | 14.9 | 16.0 | 18.2 | 3.3 | 2.2 | - |
| 4 | 16.6 | 17.3 | 18.1 | 1.5 | 0.8 | - |
| 5 | 14.9 | 16.0 | 19.2 | 4.3 | 3.2 | - |
| 6 | 18.3 | 18.3 | 19.3 | 1.0 | 1.0 | + |
| 7 | 15.4 | 16.7 | 19.3 | 3.9 | 2.6 | + |
| 8 | 18.0 | 18.6 | 19.8 | 1.8 | 1.2 | + |

#### Sertoli Cell Dilution Series and Comparison with Leydig Cells

A similar validation was performed using Sertoli cells from a control patient (Sample 1), following the experimental design previously applied to K562 cells, to establish a practical working range for archived FFPE tissue. Cellular input amounts were progressively scaled from 50 cells down to single cells (Supplementary Figure 5), alongside two negative controls. Supplementary Table 2 summarizes the number of replicates, the total area of collected cells (µm²), and the cDNA yield (ng) for each group. A detailed overview of RNA-seq metrics is presented in Supplementary Table 3. Read proportion distributions were consistent with K562 cell results.

The dilution series demonstrated a progressive increase in gene count with higher cell input (Figure 3A). For comparative purposes, larger LCM-isolated areas from the same control sample were included, consisting of individual seminiferous tubules (± 200,000 µm²). Starting from an input of 30 cells, the number of detected protein-coding genes approaches the levels observed in larger tissue areas. Samples containing 30-50 cells yielded an average of 14,189 detected protein-coding genes, whereas single-cell samples captured an average of 6,490 genes. Similar gene biotype proportions were detected as in the K562 cell dilution series (Supplementary Figure 6); however, Sertoli cell samples exhibited greater variability in library complexity and reached full saturation for most, but not all samples (Supplementary Figure 7). Sample-to-sample correlation analysis revealed generally good concordance in gene expression profiles (Figure 3B). Correlation levels were lower compared to K562 cells, likely reflecting a less homogenous population in biological setting and a reduced number of detected genes, introducing more variability.

**Figure 3.**
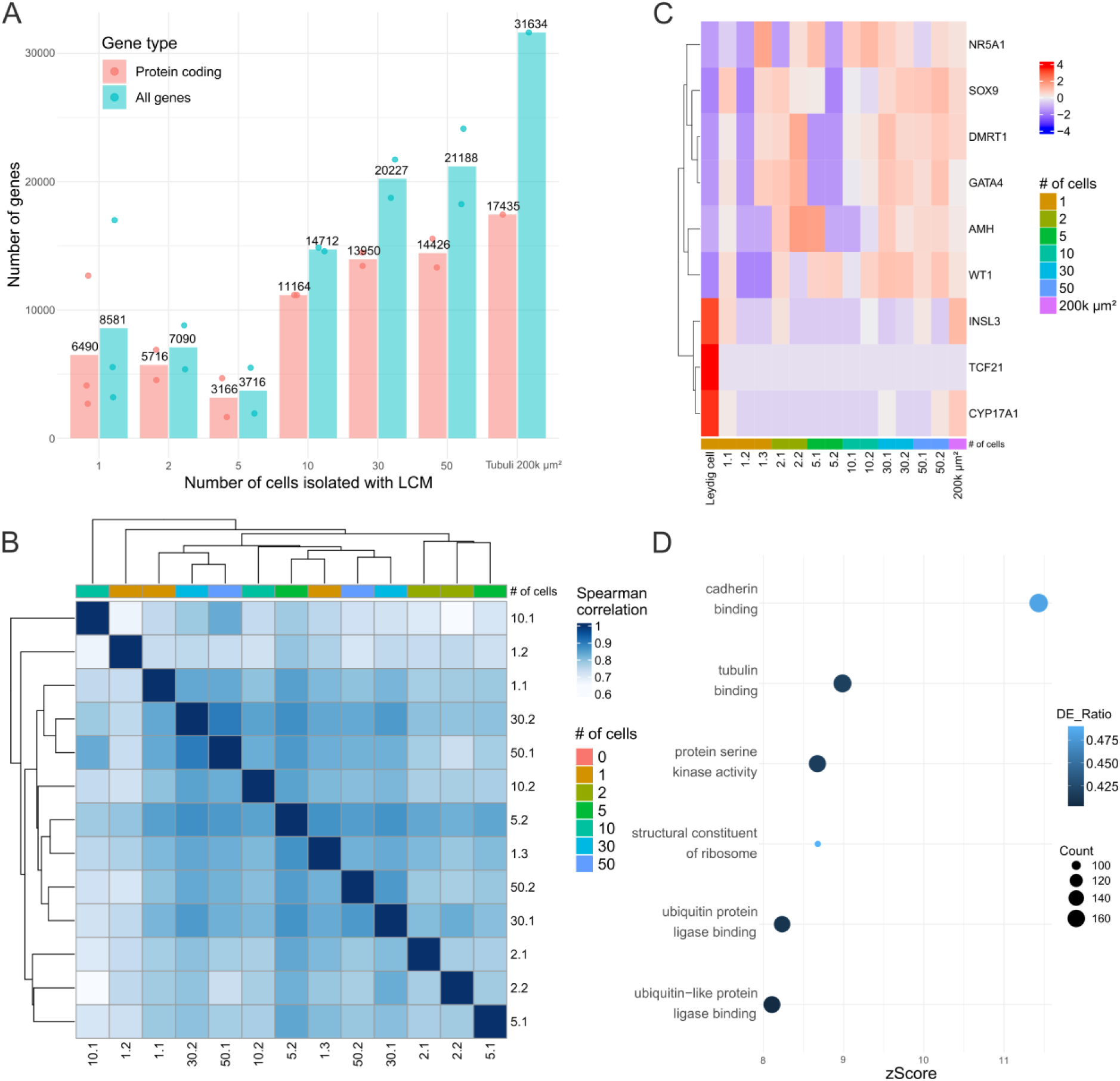
Dilution series of human Sertoli cells from a control sample, with comparative analysis of Leydig cells. (A) Average number of detected genes across varying input levels, shown separately for protein-coding genes and all detected genes. Individual replicates are shown as dots, while bars represent the mean gene count per condition. (B) Sample-to-sample correlation heatmap of expression profiles across all samples of the dilution series, based on Euclidean distance and hierarchical clustering. Sample IDs are indicated as *X.Y*, where *X* represents the number of cells isolated with LCM and *Y* indicates the replicate number. (C) Heatmap displaying expression profiles of Sertoli-specific markers (*WT1*, *AMH*, *SOX9*, *DMRT1*, *GATA4,* and *NR5A1*) and Leydig-specific markers (*TCF21*, *CYP17A1,* and *INSL3*) across the dilution series of LCM-isolated Sertoli cells. Sample IDs are indicated as *X.Y*, where *X* represents the number of cells isolated with LCM and *Y* indicates the replicate number. (D) ORA analysis for GO enrichment between Sertoli versus Leydig cells. The top terms of biological processes (BP) in GO enrichment analysis are shown. The significantly enriched pathways (adjusted p-value (P_adj_) < 0.05), derived from the GO database, are presented along with their corresponding zScores (indicating pathway enrichment direction), and the associated Gene Count (number of input genes mapping to the GO term) and DE_ratio (proportion of input genes mapping to the GO term relative to the total number of genes annotated to that GO term).

In addition to Sertoli cells, Leydig cells were isolated from the same control to verify the distinguishability of cell types using a low cell input (1-10 cells). Although substantial variance remains due to the low cell input (Supplementary Table 3), PCA demonstrated separation between Sertoli and Leydig cells (Supplementary Figure 8). To evaluate the potential contamination of other cell types during LCM, we evaluated the expression of cell-type specific markers (Figure 3C). Sertoli cell purity across the dilution series was assessed using established markers (*WT1*, *AMH*, *SOX9*, *DMRT1*, *GATA4*, and *NR5A1*), while Leydig cell identity was confirmed by the expression of *TCF21*, *CYP17A1*, and *INSL3*. The isolated Sertoli cells, particularly those from the higher-input samples (30–50 cells), exhibited strong expression of Sertoli-specific markers, with no detectable expression of Leydig-specific markers. In contrast, lower-input Sertoli samples displayed greater variability in marker expression. Leydig cell isolated from the same tissue showed clear expression of Leydig-specific markers, confirming its identity. Notably, the single-cell samples appeared to represent purer cell populations compared to the larger sample containing seminiferous tubules, where low-level expression of Leydig markers was observed, likely reflecting minor inclusion of adjacent intertubular tissue during microdissection. Comparative gene expression analysis revealed predominant upregulation of genes in Sertoli cells relative to Leydig cells (Supplementary Figure 9). Over-representation analysis (ORA) indicates a biologically consistent enrichment of cadherin-binding genes in Sertoli cells (Figure 3D). These proteins mediate cell-to-cell adhesion, a critical function of Sertoli cells in spermatogenesis and formation of the blood-testis barrier, which secures the specialized microenvironment of the seminiferous tubules (Piprek et al. 2020).

#### LCM-FFPEseq reveals GAHT-induced spermatogenic suppression and testicular fibrotic remodelling

To investigate the effects of GAHT and explore the mechanisms underlying tubular hyalinization of the seminiferous tubules, exposed testicular tissues from transgender females were compared with testicular tissues of controls. Using LCM-FFPEseq, tissue areas of 200,000 µm² were isolated, corresponding to one or two mature seminiferous tubules. Conventional bulk RNA-seq of testicular tissue would obscure seminiferous tubule-specific gene expression signals due to dilution from surrounding cell types. An average of 17,007 protein-coding genes were identified per sample (Supplementary Figure 10). PCA and hierarchical clustering clearly separated the samples into three groups: control testicular tissue, GAHT-exposed testicular tissue without tubular hyalinization, and GAHT-exposed testicular tissue with tubular hyalinization (Figure 4B-C). Differential expression analysis revealed a greater number of downregulated genes in exposed samples (3,782 downregulated vs. 873 upregulated; Figure 4D and Supplementary Figure 11), predominantly associated with spermatogenesis, with the strongest suppression observed in samples showing tubular hyalinization. In parallel, a progressive upregulation of genes related to extracellular matrix remodelling and inflammatory pathways was observed (Figure 5 and Supplementary Figure 12).

**Figure 4.**
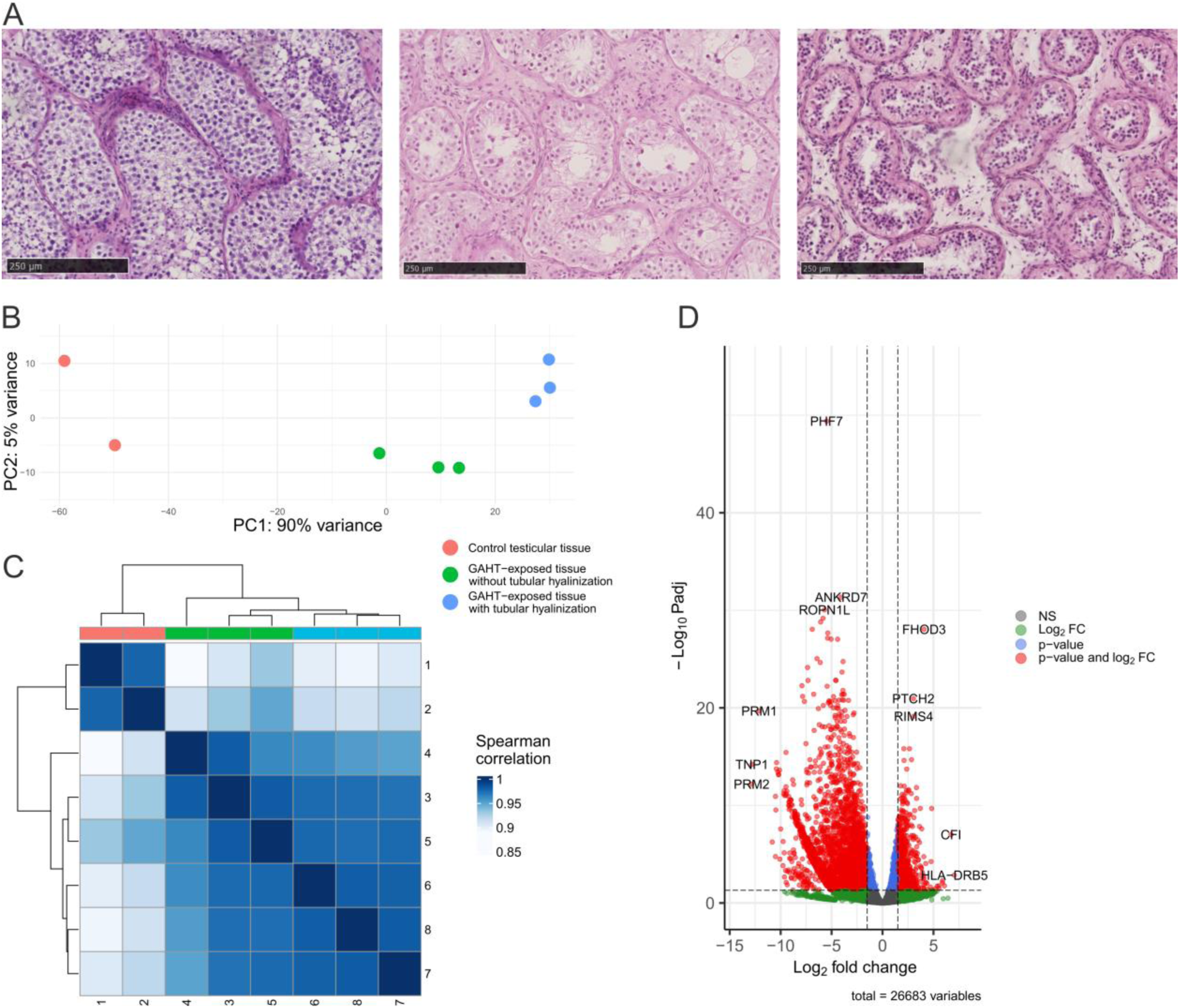
LCM-FFPEseq identifies clear separation between testicular tissue of controls, GAHT-exposed testicular tissue without and with tubular hyalinization. (A) H&E staining of included samples, from left to right: control (Sample 1), GAHT-exposed without tubular hyalinization (Sample 3), GAHT-exposed with tubular hyalinization (Sample 6). (B) PCA plot of the three different conditions. Two-dimensional PCA was used to demonstrate the clustering between control testicular tissue samples (red), GAHT-exposed testicular tissue without tubular hyalinization (green) and GAHT-exposed testicular tissue with tubular hyalinization (blue). The PCA plot was generated based on the top 500 most variable genes. (C) Sample-to-sample correlation heatmap of expression profiles across all included samples, based on Euclidean distance and hierarchical clustering. (D) Volcano plot of DEGs between GAHT-exposed testicular tissue with tubular hyalinization versus controls. The volcano plot displays log_2_FC on the x-axis and the -log10(P_adj_) on the y-axis. Single genes are depicted as dots. A total of 3,782 genes were significantly downregulated (left side) and 873 genes significantly upregulated (right side) based on the applied thresholds.

**Figure 5.**
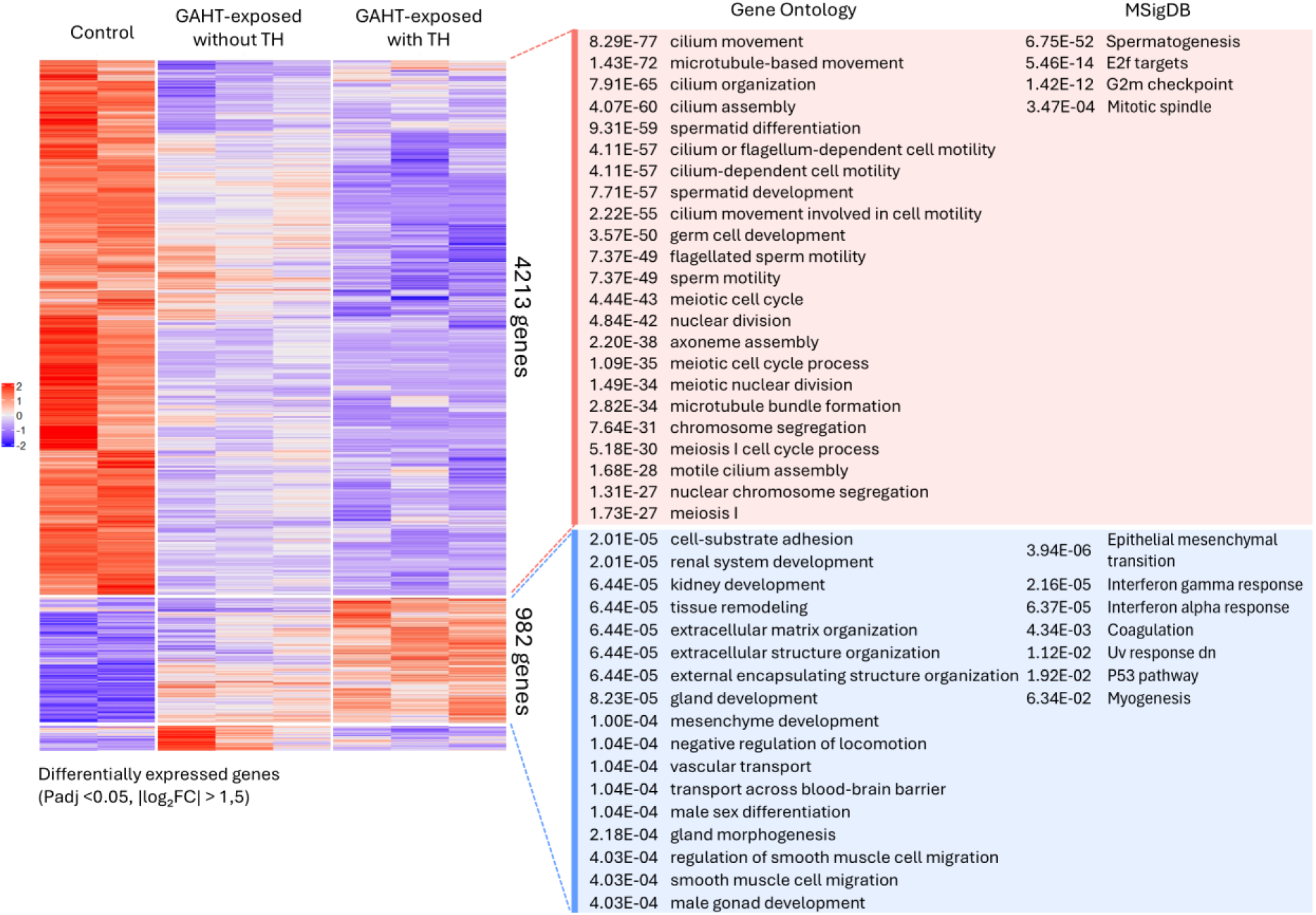
Transcriptomic alterations in testicular tissue of controls, GAHT-exposed testicular tissue without and with tubular hyalinization. Heatmap displaying 5,195 differentially expressed genes (P_adj_ < 0.05, absolute log_2_ fold change |log₂FC| > 1.5), with hierarchical clustering separating controls, therapy-exposed individuals without tubular hyalinization, and those with tubular hyalinization. Genes upregulated in the thickened group (n = 982) were enriched for Gene Ontology (GO) terms and Molecular Signatures Database (MSigDB) pathways related to extracellular matrix organization, tissue remodelling, mesenchyme development, and epithelial-mesenchymal transition, consistent with fibrotic changes. In contrast, genes downregulated in the thickened group (n = 4,213) were associated with spermatogenesis, cilium movement, meiosis, and other testis-specific processes, reflecting suppression of germ cell function and spermatogenic capacity. TH: tubular hyalinization.

## Discussion

The extensive heterogeneity of the transcriptome within tissues and cell types underscores the value of spatial resolution to uncover cell-type-specific gene expression. However, most spatial transcriptomic technologies are optimized for fresh-frozen tissue, whereas FFPE tissue remains the predominant archived format in clinical and research biobanks. Despite rapid advancements, current special transcriptomic approaches remain costly and often limited in sensitivity (Cheng et al. 2023; Guo et al. 2023). In this study, we developed LCM-FFPEseq, enabling transcriptomic analysis of LCM-isolated cells or regions of interest isolated from FFPE tissue sections, without requiring a separate RNA extraction step. By combining LCM with the latest SS3X protocol (Hagemann-Jensen et al. 2022), LCM-FFPEseq achieves single-cell resolution and surpasses existing methods in sensitivity, while its minimal input requirements streamline processing and enable transcriptomic analysis of scarce tissue or rare cell populations.

### Validation of LCM-FFPEseq

Traditional LCM-based transcriptomic workflows with separate RNA extraction steps typically require several mm² of tissue or thousands of cells to yield sufficient RNA for sequencing (Merienne et al. 2019; Paul et al. 2025). In contrast, LCM-FFPEseq demonstrated strong sensitivity across a wide input range, including single-cell conditions. In the dilution series of K562 cells, an average of 16,246 protein-coding genes were detected per sample at input levels between 30 and 500 cells. In human FFPE testicular tissue, 30-50 Sertoli cells yielded an average of 14,189 protein-coding genes. These detection rates are comparable to conventional bulk RNA-seq protocols (García-Ortega and Martínez 2015), and matched or exceeded those reported in prior LCM-based RNA-seq methods on FFPE tissue, although conducted on other cell types, such as Smart-3SEQ, which reported up to ∼10,000 genes from 500 cells (Foley et al. 2019), and LCM-seq on fresh-frozen tissue which detected ∼13,000 genes (Reads Per Kilobase of transcript, per Million mapped reads (RPKM) ≥ 1) and up to ∼17,000 genes (RPKM ≥ 0.1) from 120-cell inputs (Nichterwitz et al. 2016). Moreover, these studies do not distinguish between protein-coding and total exon gene counts, further underscoring the sensitivity of our approach. However, it should be noted that gene expressions are inherently variable and influenced by factors such as cell type, spatial context, and temporal dynamics, which complicates direct comparison across protocols (Alberts B 2002; Wang et al. 2024).

As expected, higher cell-input samples of the dilution series yielded higher cDNA quantities, confirming the efficiency of the lysis process. Starting from an input of 30 cells, protein-coding gene detection rates approach the levels of larger tissue areas with high reproducibility, sensitivity, and consistency. This cellular input is considerably lower than previous LCM-based workflows that typically require up to thousands of cells or several mm² of tissue (Merienne et al. 2019; Paul et al. 2025). In addition, LCM-FFPEseq achieved an average exon mapping rate of 34.3% for 30–500 K562 cells and 37.2% for 30-50 Sertoli cells, a significant improvement of the <10% reported in Smart-3SEQ (Foley et al. 2019), likely due to the optimizations of SS3 (Hagemann-Jensen et al. 2020) and SS3X (Hagemann-Jensen et al. 2022) which are not incorporated in Smart-3SEQ. These newer iterations incorporate crucial refinements, such as improved handling of concatemerization and strand invasion, reducing the proportion of unmapped reads and enhancing transcriptome coverage. Library complexity analysis showed that K562 samples reached saturation at ≥30 cells, while Sertoli cell samples from different patients and following medical treatments displayed more variability, indicating residual complexity and potential for higher sensitivity with deeper sequencing. Expression of established Sertoli- and Leydig-specific markers further showed consistent cell-type specificity in higher-input samples (30-50 cells). Variability at lower input levels may indicate incomplete capture or contamination of other cell types given that Histogene™ staining, while ideal for preserving RNA integrity due to its quick staining properties, lacks cell-type specificity.

Notably, LCM-FFPEseq demonstrated high sensitivity even at single-cell resolution, detecting on average 7,353 and 6,490 protein-coding genes in single K562 cells and in single Sertoli cells, respectively. These gene detection rates are in line with scRNA-seq data for these cell types (Attar et al. 2018; Natarajan et al. 2019; Zhao et al. 2020), and outperforms previous LCM-based approaches, although conducted on other cell types, such as Smart-3SEQ, which reported 5,000–6,000 genes per single FFPE cell (10), and LCM-seq, which detected ∼7,000 genes (RPKM ≥ 1) and up to ∼10,000 genes (RPKM ≥ 0.1) for single cells isolated from fresh-frozen tissue slides (9). Despite challenges in generating high-quality libraries from individual cells, LCM-FFPEseq achieved a 67% success rate for single-cell samples, increasing to 100% for inputs of five cells or more, consistent with LCM-seq’s reported performance. However, single-cell applications are constrained by the limited section thickness of FFPE tissue. This restricts the ability to capture an entire cell, particularly for large or elongated cell types like Sertoli cells and is a general limitation across all spatial transcriptomics platforms including LCM (Cheng et al. 2023; Park et al. 2023). As a result, single Sertoli cells yielded fewer protein-coding genes and showed greater variability, likely due to partial cell capture. These cells were also exposed to more laser-induced stress during isolation since they needed to be individually dissected from the surrounding cell populations, unlike the homogeneous K562 cells.

In addition to protein-coding genes, the dataset included notable numbers of lncRNAs and processed pseudogenes. The detected lncRNAs, likely polyadenylated lncRNAs captured by the poly(A)-enrichment approach, and the processed pseudogenes, likely enriched due to RNA degradation in FFPE samples, were consistent with detection rates reported for other RNA-seq library preparation protocols (Ji et al. 2024; You et al. 2021). Although long overlooked (Cheetham, Faulkner, and Dinger 2020), pseudogenes are now increasingly recognized for their widespread expression in RNA-seq datasets (Frith et al. 2006; Liao et al. 2023; Troskie et al. 2021). Notably, higher fractions of pseudogene-derived transcripts are typically observed in degraded samples due to pronounced 3’-end bias and the short fragment length (Ji et al. 2024; Raplee, Evsikov, and De Evsikova 2019). Since processed pseudogenes often retain poly(A) tails and share high sequence similarity with the 3’ regions of their parent genes, short reads are more likely to map to them, leading to their overrepresentation in degraded datasets (Raplee et al. 2019).

### GAHT-induced spermatogenic suppression and testicular fibrotic remodelling

Using LCM-FFPEseq we profiled the transcriptomic landscape of FFPE testicular tissue from transgender females who had undergone suppression of endogenous testosterone with CPA during puberty, followed by 17β-estradiol to induce bodily feminisation, and compared these profiles to controls. LCM-FFPEseq enabled targeted analysis of defined regions in the form of individual seminiferous tubules. Its high sensitivity and transcript coverage allowed detection of GAHT-associated transcriptomic alterations and their correlation with histological findings.

Distinct transcriptomic profiles were observed for GAHT-exposed testicular tissue without tubular hyalinization, and GAHT-exposed testicular tissue with tubular hyalinization, highlighting the heterogeneity of tissue responses to GAHT and the ability of LCM-FFPEseq to capture these differences. Notably, genes involved in extracellular matrix remodelling and inflammatory pathways were upregulated in GAHT-exposed samples, most pronounced in cases with tubular hyalinization. Extracellular matrix remodelling is a hallmark of testicular fibrosis, where excessive matrix proteins deposition contributes to fibrosis and impaired tissue function, as observed in conditions like cryptorchidism or chronic orchitis (Y. Xu et al. 2024). Histopathological changes including fibrosis and hyalinization following exposure to GAHT have been previously reported (Jindarak et al. 2018; Sinha, Mei, and Ferrando 2021). Our transcriptomic findings corroborate these observations. Furthermore, the upregulation of inflammatory pathways supports the hypothesis that hormonal exposure triggers a tissue injury response, potentially initiating the fibrotic remodelling. This aligns with broader models of fibrosis, where chronic injury or dysregulated signalling drives matrix deposition and scarring, ultimately compromising organ function (Bonnans, Chou, and Werb 2014).

In parallel, pathways associated with spermatogenesis were downregulated, consistent with the known suppressive effect of prolonged anti-androgen therapy on germ cell maturation (Schneider et al. 2019). These findings align with recent transcriptomic data from testicular tissue of six transgender women, which revealed a marked downregulation of genes involved in spermatogenesis, underscoring the profound inhibitory effect of GAHT on germ cell maturation, and upregulation of genes associated with inflammation and fibrosis (Delgouffe et al. 2025). No stratification based on histological features was performed. In a separate study, scRNA-seq was conducted on testicular tissue of two transgender females without accompanying histological descriptions; however, histological images provided in the study suggest that one sample lacked tubular hyalinization, while the other exhibited apparent tubular hyalinization. Similar to our findings, single-cell transcriptomic analysis revealed a more pronounced impairment of spermatogenesis in the individual with presumed tubular hyalinization (Guo et al. 2020).

### Limitations of the platform

While LCM-FFPEseq offers significant advantages for (single-cell) transcriptomic analysis from LCM-isolated cells or regions of interest from FFPE material, some limitations remain. FFPE tissue is the most widely used format for long-term sample archiving, but fixation-induced RNA degradation challenges transcriptomic analysis (Liu et al. 2022). LCM-FFPEseq addresses this by adapting the SS3X protocol, omitting tagmentation, and utilizing single-end sequencing. However, due to oligo-dT priming for poly(A) enrichment, LCM-FFPEseq displays a pronounced 3′ gene body coverage bias, as only fragments containing an intact 3′ poly(A) tail are captured. This 3′ bias is well-documented in poly(A)-selected RNA-seq protocols and commonly observed even in non-degraded, very low-input samples (Wang et al. 2016). Typically, RNA degradation is assessed post-extraction and purification by determining the RNA Integrity Number (RIN) (Schroeder et al. 2006). However, since LCM-FFPEseq omits a separate RNA extraction step, enhancing performance with minimal input, RIN values cannot be obtained. Instead, pre-amplified cDNA is evaluated on the Agilent Fragment Analyzer, with fragment length and concentration serving as indicators for the integrity of the original transcripts. In addition, LCM-FFPEseq achieves sensitivity down to the single-cell level, although this may only reflect a partial transcriptome due to partial cell capture as a consequence of section thickness, an inherent limitation of spatial transcriptomic methods (Cheng et al. 2023; Park et al. 2023). Despite its ability to handle very low input samples and reduce processing time and costs, LCM remains a labour-intensive process, limiting throughput. Future advances in LCM techniques, particularly in automation and system integration, could substantially increase efficiency. In this context, artificial intelligence holds promises for enabling rapid and automated identification of cells or regions of interest within complex tissue landscapes, thereby streamlining region of interest selection and reducing operator bias (Liu et al. 2021).

### Conclusion

LCM-FFPEseq offers a valuable and complementary approach within the rapidly evolving field of spatial transcriptomics. Unlike many current spatial platforms, which are limited by the capture area (e.g., Visium) (Cheng et al. 2023; Visium Spatial Gene Expression Reagent Kits 2024) or by targeted gene panels (e.g., MERFISH) (Cheng et al. 2023; Xia et al. 2019), LCM-FFPEseq provides whole-transcriptome coverage and allows precise isolation of morphologically defined cells or regions of interest. This makes it particularly interesting for low-input or single-cell contexts, where it captures more genes than many spatial alternatives. In addition, LCM-FFPEseq enables analysis from archived FFPE tissue, which remains incompatible with the majority of current spatial transcriptomic platforms requiring fresh-frozen tissue (Cheng et al. 2023). This utility was demonstrated on GAHT-exposed testicular tissue from transgender women which revealed transcriptional signatures of inflammation, fibrosis, and impaired spermatogenesis which could be linked to histological changes. Beyond its technical advantages, LCM-FFPEseq complements both bulk- and scRNA-seq. It enables targeted enrichment of specific tissue regions to support the interpretation of bulk RNA-seq data and provides spatial anchoring for validating scRNA-seq findings, particularly for rare or dissociation-sensitive cell types. In biological settings, single-cell applications of LCM-FFPEseq are best suited to cell types that are morphologically distinct and well-separated from surrounding tissue.

In summary, LCM-FFPEseq allows the ability to capture gene expression profiles from rare or specific cells isolated from FFPE tissues via LCM with sensitivity up to the single-cell level. This method could serve as a powerful tool to unlock FFPE pathology archives in the future, revealing cell-to-cell heterogeneity in a variety of applications, paving the way for new insights into complex tissue biology and disease mechanisms.

## Methods

### RNase-free handling

All workbenches and equipment were cleaned with RNase AWAY™ (Thermo Fisher Scientific). All steps were carried out on ice unless otherwise specified. Throughout the experiments, nuclease-free water (VWR) was exclusively utilized. Additionally, only plates and tubes certified as nuclease-free were employed.

### K562 FFPE-embedded samples

K562 tissue slides were purchased from AMSBIO (Product Code 3070-0120-5pk) and mounted on PEN membrane 1.0 NF glass slides (ZEISS). To enhance tissue adhesion, the PEN membrane glass slides were irradiated with ultraviolet (UV) light at 254 nm for 30 minutes prior to use. Finally, the slides were incubated overnight at 56°C to further improve tissue adhesion to the PEN membrane.

### Human FFPE testicular tissue samples and ethical approval

Six FFPE testicular tissue samples from transgender females receiving CPA and 17β-estradiol were selected, matched for Johnsen score. Three samples were selected with tubular hyalinization and three samples without tubular hyalinization. Control testicular tissue was obtained from diagnostic biopsy samples collected in the context of fertility preservation prior to gonadotoxic treatment for malignancy. All samples were fixed in 10% neutral buffered formalin (Sigma Aldrich), embedded in paraffin (VWR), and archived at room temperature in the Department of Pathology, Ghent University Hospital. For LCM, tissue sections were cut at 7 µm thickness and mounted on PEN membrane glass slides (Life Technologies). Finally, the slides were incubated overnight at 56°C to further improve tissue adhesion to the PEN membrane. This study was approved by the medical ethics committee of Ghent University Hospital (ONZ-2024-0029).

### Histogene™ staining

Slides were subjected to quick histological (Nissl) staining based on the Arcturus Histogene™ Staining kit (Thermo Fisher Scientific). Ethanol solutions of different concentrations for the staining protocol were prepared from ≥99.5% ethanol (VWR) and nuclease-free water.

After overnight incubation, slides were immersed 30 seconds in xylene (VWR, two times), 10 seconds in 94% ethanol (one time), 10 seconds in 85% ethanol (one time), 10 seconds in 70% ethanol (one time). Slides were then covered with Histogene™ Staining solution (Thermo Fisher Scientific) for 30 seconds. Subsequently, slides were dehydrated in rising ethanol concentrations. Slides were immersed 10 seconds in 70% ethanol (two times), 10 seconds in 80% ethanol (one time) and 10 seconds in 94% ethanol (one time). Finally, air-drying was performed shortly.

### Laser Capture Microdissection

Slides were subjected to LCM immediately after staining. Cells were dissected using the PALM Microbeam LCM system (ZEISS). Protocols of the manufacturer were followed. Cells were cut at 20x magnification while keeping laser power to a minimum. For the single cells of testicular tissue, 40x magnification was used. Following the microdissection process, cells were captured into 0.2 mL adhesive cap PCR tubes (ZEISS). After LCM, the cap was checked on its expected input using the CapCheck function. Hereafter, the cap was labelled and stored at −80°C until further library preparation steps. The whole process from staining to sample freezing never took longer than 1.5 hours for any of the samples.

### Library preparation

9 µL of lysis buffer was added to each 0.2 mL adhesive cap PCR tube, containing 5% PEG 8000 (Sigma-Aldrich), 0.1% Triton X-100 (Sigma-Aldrich), 0.125 µM LCM-FFPEseq-specific oligo-dT primer (5’-Biotin-GTCTCGTGGGCTCGGAGATGTGTATAAGAGACAGTTTTTTTTTTTTTTTTTTTTTTTTTTTTTTVN-3’, IDT), 0.5 mM dNTPs/each (Thermo Fisher Scientific), 0.5 U µL^−1^ Recombinant RNase Inhibitor (Takara) and 0.025 µg µL^−1^ Proteinase K (NEB). Volumes of PEG 8000, the oligo-dT primer, and dNTPs each were adjusted to the RT volume, consistent with SS3X protocol specifications. After dispensing, the PCR tubes are centrifuged and subjected to a denaturation step of 50 minutes at 52°C followed by a proteinase K-heat inactivation step of 10 minutes at 72°C, followed by a hold at 4°C.

Immediately following denaturation, 3 µL of the RT mix, composed of 25 mM TrisHCl pH 8.4 (Alfa Aesar), 30 mM NaCl (Thermo Fisher Scientific), 2.5 mM MgCl_2_ (Thermo Fisher Scientific), 8 mM DTT (Thermo Fisher Scientific), 1.0 mM GTP (Thermo Fisher Scientific), 0.75 µM SS3X TSO (5’-Biotin—AGAGACAGATTGCGCAATGNNNNNNNNWWrGrGrG-3’, IDT), 0.5 U µL^−1^ Recombinant RNase inhibitor (Takara), 2 U µL^−1^ Maxima H Minus reverse transcriptase (Thermo Fisher Scientific), and 0.05 mM Proteinase K inhibitor (Sigma-Aldrich) was dispensed to the tubes. The tubes were briefly centrifuged after RT mix dispensing. RT was performed at 42 °C for 90 minutes, followed by ten cycles of 50 °C for 2 minutes and 42 °C for two minutes, followed by five minutes at 85 °C and a hold at 4°C.

Next, a PCR pre-amplification was conducted, with addition of 18 µL PCR mix to each tube. The PCR mix is composed of 1X SeqAmp buffer (Takara), 0.025 U µL^−1^ SeqAmp Polymerase (Takara), 0.5 µM SS3X forward primer (5’-CTACACGACGCTCTTCCGATCT-3’, IDT) and 0.1 µM LCM-FFPEseq-specific reverse primer (5’-GTCTCGTGGGCTCGGAGAT*G*T-3’, IDT). PCR pre-amplification was performed as followed: initial denaturation at 95 °C for one minute, followed by 21 cycles of 98 °C for 10 seconds, 65 °C for 30 seconds and 68°C for four minutes. The final elongation step was conducted at 72°C for 10 minutes. The number of PCR cycles for pre-amplification can be adapted based on the input and should be determined for each project.

Subsequently, all tubes were transferred to a 96 well-plate and purified using AMPure XP beads at a 0.7X ratio. Elution was performed in 12 µL of nuclease-free H_2_O. Quality and quantity of the cDNA libraries were checked with the Fragment Analyzer (Agilent) using the High Sensitivity NGS Fragment Analysis kit (1 bp – 6000 bp, Agilent) and a separate measurement of the concentration was performed using Qubit™ dsDNA Quantification Assay Kit (Invitrogen) following the manufacturer’s protocols.

1 µL of cDNA was transferred to a new 96 well-plate for sample index PCR. To each well of this 96-well plate, 11 µL of Sample Index PCR mix was added, comprising 1X Phusion HF Buffer (Thermo Fisher Scientific), 0.2 mM dNTPs/each (Thermo Fisher Scientific), and 0.010 U µL^−1^ Phusion HF Polymerase (Thermo Fisher Scientific). Finally, 8 µL of 0.2 µM custom premixed Nextera S50X/N70 index primers, each containing 10-bp dual indexes, were added to achieve a final reaction volume of 20 µL per well. The plate was incubated for 30 seconds at 98°C, followed by 10 cycles of 10 seconds at 98°C, 30 seconds at 55°C and one minute at 72°C. The final elongation step was conducted at 72°C for five minutes. Purification of the final cDNA libraries was performed using AMPure XP beads at a 0.7X ratio with elution in 20 µL nuclease-free water.

A strict size selection was performed to remove adaptors from the final RNA-seq libraries. cDNA fragments shorter than 220 bp should be removed using E-GEL 2% (Thermo Fisher Scientific). Next, the ZYMOclean™ Gel DNA Recovery kit (ZYMO) was utilized to purify cDNA from the agarose gel with final elution in 13 µL nuclease-free water. The concentrations of the libraries were quantified using qPCR-based KAPA Library Quantification kit (qPCR) to enable equimolar pooling.

### Protocols.io

A full and comprehensive protocol of LCM-FFPEseq has been deposited on protocols.io (Reviewers link: https://www.protocols.io/private/777B6AF042CD11F0AB870A58A9FEAC02).

### Sequencing

Libraries were sequenced on the AVITI™ sequencer using a high output kit, reading 150 nt for read 1, 10 nt for P7 index read and 10 nt for P5 index read. Sequencing was performed with a target depth of approximately 15 million SE 150-bp reads per sample.

### Primary data processing

Pre-processing of the raw FASTQ files was conducted. Poly-A-trimming (cutadapt -a AAAAAAAAAA -m 20) and adapter removal (filtering pattern ‘ATTGCGCAATG([AGCT]{3})(AAAAAAAAAA)’) was performed on read 1 sequences. File synchronization was performed to maintain consistency between the filtered read 1 sequences and their corresponding index files (I1 and I2). Filtered data was then processed using zUMIs version 2.9.7e (Parekh et al. 2018). Reads were filtered for low-quality barcodes and UMIs (4 bases < phred 20, 3 bases < phred 20, respectively) and UMI-containing reads parsed by detection of the pattern (ATTGCGCAATG) while allowing up to two mismatches. Finally, within the zUMIs pipeline, filtered reads were mapped to the human reference genome (GRCh38 release 111) using STAR version 2.7.11b. with the additional parameter ‘--clip3pAdapterSeq CTGTCTCTTATACACATCT’.

### Read mapping distribution

To evaluate the read mapping distribution, read count statistics were extracted from the zUMIs output files. Three raw data tables were imported in R: total gene counts (`genecounts.txt`), UMI counts (`UMIcounts.txt`), and reads per cell (`readspercell.txt`). Sample barcodes were mapped using a separate barcode annotation file (`bc.txt`) and merged with the corresponding tables to assign sample IDs. A boxplot representing the distribution of total reads assigned to each genomic feature (ambiguity, exon, intergenic, intron and unmapped) was generated using the *ggplot2* package (Wickham 2009).

### Protein-coding gene detection and gene biotype analysis

Gene biotype annotation was performed on the filtered UMI count table derived from zUMIs output. Gene annotation data was imported from the GTF file using the *rtracklayer* package (Lawrence, Gentleman, and Carey 2009), and filtered to retain only gene-level entries. Gene annotation data was matched to the UMI count matrix. Detected genes (counts > 0) were quantified per sample and biotype using *dplyr* (Wickman H 2022) and *tidyr* (Wickham H 2024) package. To determine the number of detected protein-coding genes, only genes annotated as protein_coding in the GTF file were retained. Within each input group, the mean, minimum, and maximum number of detected genes was calculated using *dplyr* (Wickman H 2022). The resulting values were visualized using *ggplot2* (Wickham 2009) as grouped bar plots, with vertical error bars representing the observed minimum and maximum gene counts across replicates. The percentage contribution of each biotype per sample was calculated as (gene count for specific biotype / total gene count for that sample) × 100%. The average percentage contribution was then calculated per sample group, and gene biotype visualizations were created using *ggplot2* (Wickham 2009), filtering out biotype-sample pairs with <1% contribution and renormalizing remaining percentages so each sample totalled 100%.

### Filtering and normalization

zUMIs generates three UMI and three read count tables: one for exon, one for intron, and one for exon+intron counts. We chose for the inclusion of intron reads to the traditional exon counts to increase the number of detected genes and improve the resolution of the gene expression profiles, as was recommended for zUMIs (Parekh et al. 2018) and Cell Ranger of 10X Genomics (Why should I include introns for my single cell whole transcriptome Gene Expression data analysis? – 10X Genomics n.d.). The count table was further analysed using R (version 4.3.3). Raw counts were initially pre-filtered to retain only genes with a minimum of seven counts in the K562 validation samples and four counts in the human tissue samples, each present in at least two samples. These thresholds correspond to the inflection point of the gene detection curve and reflect the size of the smallest experimental group. Following pre-filtering, differential expression analysis was conducted using *DESeq2* (version 1.42.1) (Love, Huber, and Anders 2014). Normalization of raw counts was carried out using the default method, and adaptive shrinkage was applied to the log_2_FC using the ashr method in *DESeq2* (Stephens 2016).

### Clustering analysis

PCA analysis was performed on rlog-transformed count data using both the plotPCA and prcomp functions from the *DESeq2* package (Love et al. 2014) to assess variance structure. Additionally, sample–sample correlation analysis was visualized using Euclidean distance-based heatmaps with the *pheatmap* package (Kolde R 2018), annotated by cell input group.

### Gene body coverage assessment

The 5’ to 3’ gene body coverage was evaluated for the LCM-FFPEseq K562 samples of the dilution series using RSEQC (version 2.6.4) (Wang, Wang, and Li 2012). Specifically, the geneBody_coverage.py script from the RSeQC package (Wang et al. 2012) was used to analyse the distribution of the sequencing reads across the length of gene bodies. The filtered.Aligned.GeneTagged.UBcorrected.sorted.bam file, generated by zUMIs, was used as input.

### Saturation curve analysis

Sequencing saturation curves were generated using the *preseq* package (Daley and Smith 2013) and visualized with *ggplot2* (Wickham 2009). UMI count data were extracted from the dgecounts.rds file generated by the zUMIs pipeline. Sample barcodes were matched to the count data, and the resulting count matrix was exported as a .csv file for input into *preseq*. Cell input categories were analysed across replicates to assess the relationship between the total number of reads and the number of distinct reads.

### Differential gene expression analysis

Differentially expressed genes were visualized using volcano plots generated with the *EnhancedVolcano* package (Blighe K n.d.). Genes with P_adj_ < 0.05 and |log₂FC| > 1.5 were considered significantly expressed and were used for both heatmap visualization and over-representation analysis. The heatmap was generated using all differentially expressed genes from the following comparisons: (1) control testicular tissue vs. GAHT-exposed testicular tissue without tubular hyalinization, (2) control testicular tissue vs. GAHT-exposed testicular tissue with tubular hyalinization, and (3) GAHT-exposed testicular tissue without tubular hyalinization vs. GAHT-exposed testicular tissue with tubular hyalinization, using the *ComplexHeatmap* package (Gu 2022; Gu, Eils, and Schlesner 2016). Genes and samples were grouped into 3 clusters by hierarchical clustering. To visualize the expression dynamics of these gene clusters across samples, a bar chart was generated with *ggplot2 (Wickham et al. 2007)*, showing for each sample the log₂ ratio of mean expression in each gene cluster to the mean expression of the corresponding cluster in control samples. Over-representation analysis was performed using GO terms and the MSigDB (Aleksander et al. 2023; Ashburner et al. 2000; Liberzon et al. 2015; Subramanian et al. 2005), with the *clusterProfiler* package (Wu et al. 2021; S. Xu et al. 2024; Yu 2024; Yu et al. 2012). Dot plot visualization was generated with *ggplot2 (Wickham et al. 2007)*, displaying the gene count (i.e., the number of differentially expressed genes within a gene set), the DE_ratio (the ratio of this count to the total number of genes in the respective gene set), and the Z-score, representing the deviation of observed gene set overlap from the expected value under a hypergeometric distribution.

## Data Access

All raw and processed sequencing data generated in this study will be submitted to NCBI Gene Expression Omnibus. A full and comprehensive protocol of LCM-FFPEseq will be deposited on protocols.io.

## Acknowledgements

We thank Sarah De Keulenaer, Ellen De Meester, and Sylvie Decraene from the Ghent University NXTGNT sequencing core facility for their expertise and assistance with the Element AVITI sequencing runs.

## Author Contributions

**E.C.** Conceptualization, Methodology, Validation, Formal Analysis, Investigation, Data Curation, Writing – Original Draft, Visualization; **H.S.** Conceptualization, Methodology, Validation, Formal Analysis, Investigation, Data Curation, Writing – Original Draft, Visualization; **K.D.** Methodology, Writing – Review & Editing; **G.C.** Methodology, Writing – Review & Editing; **H.C.** Investigation, Writing – Review & Editing; **J.V.D.V.** Investigation, Writing – Review & Editing; **D.B.** Methodology, Writing – Review & Editing; **A.V.** Methodology, Writing – Review & Editing; **D.D.** Resources, Writing - Review & Editing, Funding Acquisition; **W.D.S.** Methodology, Resources, Writing – Review & Editing; **K.V.V.** Resources, Writing – Review & Editing; **M.C.** Conceptualization, Resources, Writing – Review & Editing, Supervision, Funding Acquisition; **F.V.N.** Conceptualization, Resources, Writing – Review & Editing, Supervision, Funding Acquisition

## References

Alberts B, Johnson A, Lewis J, et al. 2002. “An Overview of Gene Control.” in Molecular Biology of the Cell. 4th edition. New York: Garland Science.

Aleksander, Suzi A., James Balhoff, Seth Carbon, J. Michael Cherry, Harold J. Drabkin, Dustin Ebert, Marc Feuermann, Pascale Gaudet, Nomi L. Harris, David P. Hill, and, et al. 2023. “The Gene Ontology Knowledgebase in 2023.” Genetics 224(1). doi:10.1093/GENETICS/IYAD031.

Ashburner, Michael, Catherine A. Ball, Judith A. Blake, David Botstein, Heather Butler, J. Michael Cherry, Allan P. Davis, Kara Dolinski, Selina S. Dwight, Janan T. Eppig, Midori A. Harris, David P. Hill, Laurie Issel-Tarver, Andrew Kasarskis, Suzanna Lewis, John C. Matese, Joel E. Richardson, Martin Ringwald, Gerald M. Rubin, and Gavin Sherlock. 2000. “Gene Ontology: Tool for the Unification of Biology.” Nature Genetics 25(1):25–29. doi:10.1038/75556;KWRD=BIOMEDICINE.

Asslaber, M., and K. Zatloukal. 2007. “Biobanks: Transnational, European and Global Networks.” Briefings in Functional Genomics and Proteomics 6(3):193–201. doi:10.1093/bfgp/elm023.

Attar, Moustafa, Eshita Sharma, Shuqiang Li, Claire Bryer, Laura Cubitt, John Broxholme, Helen Lockstone, James Kinchen, Alison Simmons, Paolo Piazza, David Buck, Kenneth J. Livak, and Rory Bowden. 2018. “A Practical Solution for Preserving Single Cells for RNA Sequencing.” Scientific Reports 2018 8:1 8(1):1–10. doi:10.1038/s41598-018-20372-7.

Blighe K, Rana S, Lewis M. n.d. “EnhancedVolcano: Publication-Ready Volcano Plots with Enhanced Colouring and Labeling.” 2025.

Bonnans, Caroline, Jonathan Chou, and Zena Werb. 2014. “Remodelling the Extracellular Matrix in Development and Disease.” Nature Reviews. Molecular Cell Biology 15(12):786–801. doi:10.1038/nrm3904.

Cheetham, Seth W., Geoffrey J. Faulkner, and Marcel E. Dinger. 2020. “Overcoming Challenges and Dogmas to Understand the Functions of Pseudogenes.” Nature Reviews Genetics 21(3):191–201. doi:10.1038/s41576-019-0196-1.

Chen, Jun, Shengbao Suo, Patrick Pl Tam, Jing Dong J. Han, Guangdun Peng, and Naihe Jing. 2017. “Spatial Transcriptomic Analysis of Cryosectioned Tissue Samples with Geo-Seq.” Nature Protocols 12(3):566–80. doi:10.1038/NPROT.2017.003.

Cheng, Mengnan, Yujia Jiang, Jiangshan Xu, Alexios-Fotios A. Mentis, Shuai Wang, Huiwen Zheng, Sunil Kumar Sahu, Longqi Liu, and Xun Xu. 2023. “Spatially Resolved Transcriptomics: A Comprehensive Review of Their Technological Advances, Applications, and Challenges.” Journal of Genetics and Genomics 50(9):625–40. doi:10.1016/j.jgg.2023.03.011.

Clément-Ziza, Mathieu, Arnold Munnich, Stanislas Lyonnet, Francis Jaubert, and Claude Besmond. 2008. “Stabilization of RNA during Laser Capture Microdissection by Performing Experiments under Argon Atmosphere or Using Ethanol as a Solvent in Staining Solutions.” RNA 14(12):2698–2704. doi:10.1261/rna.1261708.

Coleman, E., A. E. Radix, W. P. Bouman, G. R. Brown, A. L. C. de Vries, M. B. Deutsch, R. Ettner, L. Fraser, M. Goodman, J. Green, and et al. 2022. “Standards of Care for the Health of Transgender and Gender Diverse People, Version 8.” International Journal of Transgender Health 23(Suppl 1):S1–259. doi:10.1080/26895269.2022.2100644.

Cornejo, Kristine M., Esther Oliva, Rory Crotty, Peter M. Sadow, Kyle Devins, Anton Wintner, and Chin-Lee Wu. 2022. “Clinicopathologic Features and Proposed Grossing Protocol of Orchiectomy Specimens Performed for Gender Affirmation Surgery.” Human Pathology 127:21–27. doi:10.1016/j.humpath.2022.05.017.

Daley, Timothy, and Andrew D. Smith. 2013. “Predicting the Molecular Complexity of Sequencing Libraries.” Nature Methods 10(4):325–27. doi:10.1038/nmeth.2375.

Datta, Soma, Lavina Malhotra, Ryan Dickerson, Scott Chaffee, Chandan K. Sen, and Sashwati Roy. 2015. “Laser Capture Microdissection: Big Data from Small Samples.” Histology and Histopathology 30(11):1255–69.

Delgouffe, Emily, Samuel Madureira Silva, Frédéric Chalmel, Wilfried Cools, Camille Raets, Kelly Tilleman, Guy T’Sjoen, Yoni Baert, and Ellen Goossens. 2025. “Partial Rejuvenation of the Spermatogonial Stem Cell Niche after Gender-Affirming Hormone Therapy in Trans Women.” ELife 13. doi:10.7554/ELIFE.94825.

Emmert-Buck, Michael R., Robert F. Bonner, Paul D. Smith, Rodrigo F. Chuaqui, Zhengping Zhuang, Seth R. Goldstein, Rhonda A. Weiss, and Lance A. Liotta. 1996. “Laser Capture Microdissection.” Science 274(5289):998–1001. doi:10.1126/science.274.5289.998.

Evers, David L., Junkun He, Yeon Ho Kim, Jeffrey T. Mason, and Timothy J. O’Leary. 2011. “Paraffin Embedding Contributes to RNA Aggregation, Reduced RNA Yield, and Low RNA Quality.” The Journal of Molecular Diagnostics : JMD 13(6):687–94. doi:10.1016/j.jmoldx.2011.06.007.

Foley, Joseph W., Chunfang Zhu, Philippe Jolivet, Shirley X. Zhu, Peipei Lu, Michael J. Meaney, and Robert B. West. 2019. “Gene Expression Profiling of Single Cells from Archival Tissue with Laser-Capture Microdissection and Smart-3SEQ.” Genome Research 29(11):1816–25. doi:10.1101/GR.234807.118/-/DC1.

Frith, Martin C., Laurens G. Wilming, Alistair Forrest, Hideya Kawaji, Sin Lam Tan, Claes Wahlestedt, Vladimir B. Bajic, Chikatoshi Kai, Jun Kawai, Piero Carninci, Yoshihide Hayashizaki, Timothy L. Bailey, and Lukasz Huminiecki. 2006. “Pseudo–Messenger RNA: Phantoms of the Transcriptome.” PLoS Genetics 2(4):e23. doi:10.1371/journal.pgen.0020023.

García-Ortega, Luis Fernando, and Octavio Martínez. 2015. “How Many Genes Are Expressed in a Transcriptome? Estimation and Results for RNA-Seq.” PLoS ONE 10(6):e0130262. doi:10.1371/JOURNAL.PONE.0130262.

Gu, Zuguang. 2022. “Complex Heatmap Visualization.” IMeta 1(3):e43. doi:10.1002/IMT2.43;PAGEGROUP:STRING:PUBLICATION.

Gu, Zuguang, Roland Eils, and Matthias Schlesner. 2016. “Complex Heatmaps Reveal Patterns and Correlations in Multidimensional Genomic Data.” Bioinformatics 32(18):2847–49. doi:10.1093/BIOINFORMATICS/BTW313.

Guo, Jingtao, Xichen Nie, Maria Giebler, Hana Mlcochova, Yueqi Wang, Edward J. Grow, Robin Kim, Melissa Tharmalingam, Gabriele Matilionyte, Cecilia Lindskog, Douglas T. Carrell, Rod T. Mitchell, Anne Goriely, James M. Hotaling, and Bradley R. Cairns. 2020. “The Dynamic Transcriptional Cell Atlas of Testis Development during Human Puberty.” Cell Stem Cell 26(2):262–276.e4. doi:10.1016/J.STEM.2019.12.005.

Guo, Wenbo, Yining Hu, Jingyang Qian, Lidan Zhu, Junyun Cheng, Jie Liao, and Xiaohui Fan. 2023. “Laser Capture Microdissection for Biomedical Research: Towards High-Throughput, Multi-Omics, and Single-Cell Resolution.” Journal of Genetics and Genomics 50(9):641–51. doi:10.1016/j.jgg.2023.07.011.

Hagemann-Jensen, Michael, Christoph Ziegenhain, Ping Chen, Daniel Ramsköld, Gert Jan Hendriks, Anton J. M. Larsson, Omid R. Faridani, and Rickard Sandberg. 2020. “Single-Cell RNA Counting at Allele and Isoform Resolution Using Smart-Seq3.” Nature Biotechnology 2020 38:6 38(6):708–14. doi:10.1038/s41587-020-0497-0.

Hagemann-Jensen, Michael, Christoph Ziegenhain, and Rickard Sandberg. 2022. “Scalable Single-Cell RNA Sequencing from Full Transcripts with Smart-Seq3xpress.” Nature Biotechnology 2022 40:10 40(10):1452–57. doi:10.1038/s41587-022-01311-4.

Ji, Bingqing, Jiale Chen, Hui Gong, and Xiangning Li. 2024. “Streamlined Full-Length Total RNA Sequencing of Paraformaldehyde-Fixed Brain Tissues.” International Journal of Molecular Sciences 25(12):6504. doi:10.3390/ijms25126504.

Jin, Yang, Yuanli Zuo, Gang Li, Wenrong Liu, Yitong Pan, Ting Fan, Xin Fu, Xiaojun Yao, and Yong Peng. 2024. “Advances in Spatial Transcriptomics and Its Applications in Cancer Research.” Molecular Cancer 23(1):129. doi:10.1186/s12943-024-02040-9.

Jindarak, Sirachai, Kasama Nilprapha, Taywin Atikankul, Apichai Angspatt, Pornthep Pungrasmi, Seree Iamphongsai, Pasu Promniyom, Poonpissamai Suwajo, Gennaro Selvaggi, and Preecha Tiewtranon. 2018. “Spermatogenesis Abnormalities Following Hormonal Therapy in Transwomen.” BioMed Research International 2018(1):7919481. doi:10.1155/2018/7919481.

Kolde R. 2018. “Pheatmap: Pretty Heatmaps.”

Lawrence, Michael, Robert Gentleman, and Vincent Carey. 2009. “Rtracklayer: An R Package for Interfacing with Genome Browsers.” Bioinformatics 25(14):1841–42. doi:10.1093/bioinformatics/btp328.

Li, Xinmin, and Cun-Yu Wang. 2021. “From Bulk, Single-Cell to Spatial RNA Sequencing.” International Journal of Oral Science 13(1):36. doi:10.1038/s41368-021-00146-0.

Liao, Yuhan, Zhenyu Liu, Yu Zhang, Ping Lu, Lu Wen, and Fuchou Tang. 2023. “High-Throughput and High-Sensitivity Full-Length Single-Cell RNA-Seq Analysis on Third-Generation Sequencing Platform.” Cell Discovery 9(1):5. doi:10.1038/s41421-022-00500-4.

Liberzon, Arthur, Chet Birger, Helga Thorvaldsdóttir, Mahmoud Ghandi, Jill P. Mesirov, and Pablo Tamayo. 2015. “The Molecular Signatures Database Hallmark Gene Set Collection.” Cell Systems 1(6):417–25. doi:10.1016/J.CELS.2015.12.004.

Liu, Jianfei, Christine Shen, Nancy Aguilera, Catherine Cukras, Robert B. Hufnagel, Wadih M. Zein, Tao Liu, and Johnny Tam. 2021. “Active Cell Appearance Model Induced Generative Adversarial Networks for Annotation-Efficient Cell Segmentation and Identification on Adaptive Optics Retinal Images.” IEEE Transactions on Medical Imaging 40(10):2820–31. doi:10.1109/TMI.2021.3055483.

Liu, Yuanhang, Aditya Bhagwate, Stacey J. Winham, Melissa T. Stephens, Brent W. Harker, Samantha J. McDonough, Melody L. Stallings-Mann, Ethan P. Heinzen, Robert A. Vierkant, Tanya L. Hoskin, Marlene H. Frost, Jodi M. Carter, Michael E. Pfrender, Laurie Littlepage, Derek C. Radisky, Julie M. Cunningham, Amy C. Degnim, and Chen Wang. 2022. “Quality Control Recommendations for RNASeq Using FFPE Samples Based on Pre-Sequencing Lab Metrics and Post-Sequencing Bioinformatics Metrics.” BMC Medical Genomics 15(1):1–12. doi:10.1186/S12920-022-01355-0/FIGURES/4.

Love, Michael I., Wolfgang Huber, and Simon Anders. 2014. “Moderated Estimation of Fold Change and Dispersion for RNA-Seq Data with DESeq2.” Genome Biology 15(12):550. doi:10.1186/s13059-014-0550-8.

Merienne, Nicolas, Cécile Meunier, Anne Schneider, Jonathan Seguin, Satish S. Nair, Anne B. Rocher, Stéphanie Le Gras, Céline Keime, Richard Faull, Luc Pellerin, Jean-Yves Chatton, Christian Neri, Karine Merienne, and Nicole Déglon. 2019. “Cell-Type-Specific Gene Expression Profiling in Adult Mouse Brain Reveals Normal and Disease-State Signatures.” Cell Reports 26(9):2477–2493.e9. doi:10.1016/j.celrep.2019.02.003.

Natarajan, Kedar Nath, Zhichao Miao, Miaomiao Jiang, Xiaoyun Huang, Hongpo Zhou, Jiarui Xie, Chunqing Wang, Shishang Qin, Zhikun Zhao, Liang Wu, Naibo Yang, Bo Li, Yong Hou, Shiping Liu, and Sarah A. Teichmann. 2019. “Comparative Analysis of Sequencing Technologies for Single-Cell Transcriptomics.” Genome Biology 20(1):1–8. doi:10.1186/S13059-019-1676-5/FIGURES/2.

Nichterwitz, Susanne, Geng Chen, Julio Aguila Benitez, Marlene Yilmaz, Helena Storvall, Ming Cao, Rickard Sandberg, Qiaolin Deng, and Eva Hedlund. 2016. “Laser Capture Microscopy Coupled with Smart-Seq2 for Precise Spatial Transcriptomic Profiling.” Nature Communications 2016 7:1 7(1):1–11. doi:10.1038/ncomms12139.

de Nie, I., C. L. Mulder, A. Meißner, Y. Schut, E. M. Holleman, W. B. van der Sluis, S. E. Hannema, M. den Heijer, J. Huirne, A. M. M. van Pelt, and N. M. van Mello. 2022. “Histological Study on the Influence of Puberty Suppression and Hormonal Treatment on Developing Germ Cells in Transgender Women.” Human Reproduction (Oxford, England) 37(2):297–308. doi:10.1093/humrep/deab240.

PALM Protocols-DNA Handling. n.d.

Parekh, Swati, Christoph Ziegenhain, Beate Vieth, Wolfgang Enard, and Ines Hellmann. 2018. “ZUMIs - A Fast and Flexible Pipeline to Process RNA Sequencing Data with UMIs.” GigaScience 7(6). doi:10.1093/gigascience/giy059.

Park, Han Eol, Song Hyun Jo, Rosalind H. Lee, Christian P. Macks, Taeyun Ku, Jihwan Park, Chung Whan Lee, Junho K. Hur, and Chang Ho Sohn. 2023. “Spatial Transcriptomics: Technical Aspects of Recent Developments and Their Applications in Neuroscience and Cancer Research.” Advanced Science 10(16):2206939. doi:10.1002/ADVS.202206939;REQUESTEDJOURNAL:JOURNAL:21983844;WGROUP:STRING:PUBLICATION.

Paul, Evan D., Barbora Huraiová, Natália Valková, Natalia Matyasovska, Daniela Gábrišová, Soňa Gubová, Helena Ignačáková, Tomáš Ondris, Michal Gala, Liliane Barroso, Silvia Bendíková, Jarmila Bíla, Katarína Buranovská, Diana Drobná, Zuzana Krchňáková, Maryna Kryvokhyzha, Daniel Lovíšek, Viktoriia Mamoilyk, Veronika Mancikova, Nina Vojtaššáková, Michaela Ristová, Iñaki Comino-Méndez, Igor Andrašina, Pavel Morozov, Thomas Tuschl, Fresia Pareja, Jakob N. Kather, and Pavol Čekan. 2025. “The Spatially Informed MFISHseq Assay Resolves Biomarker Discordance and Predicts Treatment Response in Breast Cancer.” Nature Communications 16(1):226. doi:10.1038/s41467-024-55583-2.

Picelli, Simone, Omid R. Faridani, Åsa K. Björklund, Gösta Winberg, Sven Sagasser, and Rickard Sandberg. 2014. “Full-Length RNA-Seq from Single Cells Using Smart-Seq2.” Nature Protocols 2013 9:1 9(1):171–81. doi:10.1038/nprot.2014.006.

Piprek, Rafał P., Malgorzata Kloc, Paulina Mizia, and Jacek Z. Kubiak. 2020. “The Central Role of Cadherins in Gonad Development, Reproduction, and Fertility.” International Journal of Molecular Sciences 2020, Vol. 21, Page 8264 21(21):8264. doi:10.3390/IJMS21218264.

Raplee, Isaac D., Alexei V. Evsikov, and Caralina Marín De Evsikova. 2019. “Aligning the Aligners: Comparison of RNA Sequencing Data Alignment and Gene Expression Quantification Tools for Clinical Breast Cancer Research.” Journal of Personalized Medicine 9(2):18. doi:10.3390/JPM9020018.

Schneider, Florian, Bettina Scheffer, Jennifer Dabel, Laura Heckmann, Stefan Schlatt, Sabine Kliesch, and Nina Neuhaus. 2019. “Options for Fertility Treatments for Trans Women in Germany.” Journal of Clinical Medicine 2019, Vol. 8, Page 730 8(5):730. doi:10.3390/JCM8050730.

Schroeder, Andreas, Odilo Mueller, Susanne Stocker, Ruediger Salowsky, Michael Leiber, Marcus Gassmann, Samar Lightfoot, Wolfram Menzel, Martin Granzow, and Thomas Ragg. 2006. “The RIN: An RNA Integrity Number for Assigning Integrity Values to RNA Measurements.” BMC Molecular Biology 7(1):1–14. doi:10.1186/1471-2199-7-3/FIGURES/8.

Simone, Nicole L., Robert F. Bonner, John W. Gillespie, Michael R. Emmert-Buck, and Lance A. Liotta. 1998. “Laser-Capture Microdissection: Opening the Microscopic Frontier to Molecular Analysis.” Trends in Genetics 14(7):272–76. doi:10.1016/S0168-9525(98)01489-9.

Sinha, Annika, Lin Mei, and Cecile Ferrando. 2021. “The Effect of Estrogen Therapy on Spermatogenesis in Transgender Women.” F&S Reports 2(3):347–51. doi:10.1016/J.XFRE.2021.06.002.

Stephens, Matthew. 2016. “False Discovery Rates: A New Deal.” Biostatistics kxw041. doi:10.1093/biostatistics/kxw041.

Subramanian, Aravind, Pablo Tamayo, Vamsi K. Mootha, Sayan Mukherjee, Benjamin L. Ebert, Michael A. Gillette, Amanda Paulovich, Scott L. Pomeroy, Todd R. Golub, Eric S. Lander, and Jill P. Mesirov. 2005. “Gene Set Enrichment Analysis: A Knowledge-Based Approach for Interpreting Genome-Wide Expression Profiles.” Proceedings of the National Academy of Sciences of the United States of America 102(43):15545–50. doi:10.1073/PNAS.0506580102/SUPPL_FILE/06580FIG7.JPG.

Troskie, Robin-Lee, Yohaann Jafrani, Tim R. Mercer, Adam D. Ewing, Geoffrey J. Faulkner, and Seth W. Cheetham. 2021. “Long-Read CDNA Sequencing Identifies Functional Pseudogenes in the Human Transcriptome.” Genome Biology 22(1):146. doi:10.1186/s13059-021-02369-0.

Vahrenkamp, Jeffery M., Kathryn Szczotka, Mark K. Dodson, Elke A. Jarboe, Andrew P. Soisson, and Jason Gertz. 2019. “FFPEcap-Seq: A Method for Sequencing Capped RNAs in Formalin-Fixed Paraffin-Embedded Samples.” Genome Research 29(11):1826–35. doi:10.1101/GR.249656.119/-/DC1.

Visium Spatial Gene Expression Reagent Kits. 2024. www.10xgenomics.com/trademarks.

Waldron, Levi, Peter Simpson, Giovanni Parmigiani, and Curtis Huttenhower. 2012. “Report on Emerging Technologies for Translational Bioinformatics: A Symposium on Gene Expression Profiling for Archival Tissues.” BMC Cancer 12:124. doi:10.1186/1471-2407-12-124.

Wang, Hongyang, James D. Owens, Joanna H. Shih, Ming-Chung Li, Robert F. Bonner, and J. Frederic Mushinski. 2006. “Histological Staining Methods Preparatory to Laser Capture Microdissection Significantly Affect the Integrity of the Cellular RNA.” BMC Genomics 7(1):97. doi:10.1186/1471-2164-7-97.

Wang, Liguo, Jinfu Nie, Hugues Sicotte, Ying Li, Jeanette E. Eckel-Passow, Surendra Dasari, Peter T. Vedell, Poulami Barman, Liewei Wang, Richard Weinshiboum, Jin Jen, Haojie Huang, Manish Kohli, and Jean Pierre A. Kocher. 2016. “Measure Transcript Integrity Using RNA-Seq Data.” BMC Bioinformatics 17(1):1–16. doi:10.1186/S12859-016-0922-Z/FIGURES/8.

Wang, Liguo, Shengqin Wang, and Wei Li. 2012. “RSeQC: Quality Control of RNA-Seq Experiments.” Bioinformatics 28(16):2184–85. doi:10.1093/bioinformatics/bts356.

Wang, Siying, Shichao Lin, and Chaoyong Yang. 2024. “The Dawn of Spatiotemporal Transcriptomics.” Biomedical Analysis 1(2):154–61. doi:10.1016/j.bioana.2024.06.002.

Why should I include introns for my single cell whole transcriptome Gene Expression data analysis? – 10X Genomics. n.d. Retrieved June 23, 2025. https://kb.10xgenomics.com/hc/en-us/articles/4998628924429-Why-should-I-include-introns-for-my-single-cell-whole-transcriptome-Gene-Expression-data-analysis.

Wickham H, Vaughan D, Girlich M. 2024. “Tidyr: Tidy Messy Data.”

Wickham, Hadley. 2009. Ggplot2. New York, NY: Springer New York.

Wickham, Hadley, Winston Chang, Lionel Henry, Thomas Lin Pedersen, Kohske Takahashi, Claus Wilke, Kara Woo, Hiroaki Yutani, Dewey Dunnington, and Teun van den Brand. 2007. “Ggplot2: Create Elegant Data Visualisations Using the Grammar of Graphics.” CRAN: Contributed Packages.

Wickman H, François R, Henry L, Müller K. 2022. “Dplyr: A Grammar of Data Manipulation.”

Williams, Cameron G., Hyun Jae Lee, Takahiro Asatsuma, Roser Vento-Tormo, and Ashraful Haque. 2022. “An Introduction to Spatial Transcriptomics for Biomedical Research.” Genome Medicine 14(1):68. doi:10.1186/s13073-022-01075-1.

Wu, Tianzhi, Erqiang Hu, Shuangbin Xu, Meijun Chen, Pingfan Guo, Zehan Dai, Tingze Feng, Lang Zhou, Wenli Tang, Li Zhan, Xiaocong Fu, Shanshan Liu, Xiaochen Bo, and Guangchuang Yu. 2021. “ClusterProfiler 4.0: A Universal Enrichment Tool for Interpreting Omics Data.” The Innovation 2(3):100141. doi:10.1016/j.xinn.2021.100141.

Xia, Chenglong, Hazen P. Babcock, Jeffrey R. Moffitt, and Xiaowei Zhuang. 2019. “Multiplexed Detection of RNA Using MERFISH and Branched DNA Amplification.” Scientific Reports 9(1):7721. doi:10.1038/s41598-019-43943-8.

Xu, Shuangbin, Erqiang Hu, Yantong Cai, Zijing Xie, Xiao Luo, Li Zhan, Wenli Tang, Qianwen Wang, Bingdong Liu, Rui Wang, Wenqin Xie, Tianzhi Wu, Liwei Xie, and Guangchuang Yu. 2024. “Using ClusterProfiler to Characterize Multiomics Data.” Nature Protocols 19(11):3292–3320. doi:10.1038/s41596-024-01020-z.

Xu, Ying, Poyi Hu, Wanyi Chen, Jin Chen, Chunyan Liu, and Huiping Zhang. 2024. “Testicular Fibrosis Pathology, Diagnosis, Pathogenesis, and Treatment: A Perspective on Related Diseases.” Andrology. doi:10.1111/andr.13769.

You, Yue, Luyi Tian, Shian Su, Xueyi Dong, Jafar S. Jabbari, Peter F. Hickey, and Matthew E. Ritchie. 2021. “Benchmarking UMI-Based Single-Cell RNA-Seq Preprocessing Workflows.” Genome Biology 22(1):339. doi:10.1186/s13059-021-02552-3.

Yu, Guangchuang. 2024. “Thirteen Years of ClusterProfiler.” The Innovation 5(6):100722. doi:10.1016/j.xinn.2024.100722.

Yu, Guangchuang, Li-Gen Wang, Yanyan Han, and Qing-Yu He. 2012. “ClusterProfiler: An R Package for Comparing Biological Themes Among Gene Clusters.” OMICS: A Journal of Integrative Biology 16(5):284–87. doi:10.1089/omi.2011.0118.

Zhao, Liang Yu, Chen Cheng Yao, Xiao Yu Xing, Tao Jing, Peng Li, Zi Jue Zhu, Chao Yang, Jing Zhai, Ru Hui Tian, Hui Xing Chen, Jia Qiang Luo, Na Chuan Liu, Zhi Wen Deng, Xiao Han Lin, Na Li, Jing Fang, Jie Sun, Chen Chen Wang, Zhi Zhou, and Zheng Li. 2020. “Single-Cell Analysis of Developing and Azoospermia Human Testicles Reveals Central Role of Sertoli Cells.” Nature Communications 11(1). doi:10.1038/S41467-020-19414-4.

