## Supplementary for "A laser capture microdissection-based method for high-sensitivity transcriptomics from archived FFPE tissue slides with single-cell resolution using LCM-FFPEseq"

### Supplemental Material

#### Supplementary Figures

**Supplementary Figure 1.** Detailed schematic overview of the LCM-FFPEseq library preparation protocol.

**Adapter and primer sequences:**

- LCM-FFPEseq-specific oligo-dT primer : 5'-/5Biosg/GTCTCGTGGGCTCGGAGATGTGTATAAGAGACAG(T)30VN-3'
- SS3X TSO : 5′-/5BiosG/AGAGACAGATTGCGCAATGNNNNNNNNWWrGrGrG-3′
- SS3X FWP cDNA-preamplification : 5′-TCGTCGGCAGCGTCAGATGTGTATAAGAGACAGATTGCGCAA*T*G-3′ (*=Phosphorothioated DNA bases)
- LCM-FFPEseq-specific RVP cDNA-preamplification : 5′-GTCTCGTGGGCTCGGAGAT*G*T-3′ (*=Phosphorothioated DNA bases)
- SS3X Nextera Unique Dual Index FWP : 5′-AATGATACGGCGACCACCGAGATCTACACNNNNNNNNTCGTCGGCAGCGTC-3′
- SS3X Nextera Unique Dual Index RVP : 5′-CAAGCAGAAGACGGCATACGAGAT-NNNNNNNN-GTCTCGTGGGCTCGG-3′

**Step-by-step library generation:**

1) Cell lysis: release of RNA
2) mRNA

5′ -----------------------------------$(A)_n$ 3′

3) RT

a. + oligo-dT primer

5′ ------------------------------------$(A)_n$ 3′

3′ NV$(T)_{30}$GACAGAGAATATGTGTAGAGGCTCGGGTGCTCTG 5′

b. + Maxima H Minus Reverse Transcriptase

5′ ------------------------------------$(A)_n$ 3′

3′ ← NV$(T)_{30}$GACAGAGAATATGTGTAGAGGCTCGGGTGCTCTG 5′

c. Terminal transferase activity incorporates 3 extra cytosines

5′ ------------------------------------$(A)_n$ 3′

3′ CCC-------------------------------NV$(T)_{30}$GACAGAGAATATGTGTAGAGGCTCGGGTGCTCTG 5′

d. Non-templated attachment of TSO

+ TSO: 5′ /5BiosG/AGAGACAG-ATTGCGCAATG-NNNNNNNN-WW-rGrGrG 3′
Tn5 Motif Tag UMI Spacer

5′/AGAGACAGATTGCGCAATGNNNNNNNNWWrGrGrG ----------------$(A)_n$ 3′

3′ C C C --------------NV$(T)_{30}$GACAGAGAATATGTGTAGAGGCTCGGGTGCTCTG 5′

e. Template-switching

5′ AGAGACAGATTGCGCAATGNNNNNNNNNWWrGrGrG ------------------------(A)$_{30}$CTGTCTCTTATACACATCTCCGAGCCCACGAGAC 3′

3′ TCTCTGTCTAACGCGTTACNNNNNNNNNWW CCC ------------------------NV(T)$_{30}$GACAGAGAATATGTGTAGAGGCTCGGGTGCTCTG 5′

4) PCR preamplification (x PCR cycles)

FWP: 5′ TCGTCGGCAGCGTCAGATGTGTATAAGAGACAGATTGCGCAA*T*G 3′

RVP: 5′ GTCTCGTGGGCTCGGAGAT*G*T 3′

5′AGAGACAGATTGCGCAATGNNNNNNNNNWWrGrGrG ----(A)$_{30}$CTGTCTCTTATACACATCTCCGAGCCCACGAGAC 3′
← 3′ TGTAGAGGCTCGGGTGCTCTG 5′
5′TCGTCGGCAGCGTCAGATGTGTATAAGAGACAGATTGCGCAATG 3′→
3′TCTCTGTCTAACGCGTTACNNNNNNNNNWWCCC----NV(T)$_{30}$GACAGAGAATATGTGTAGAGGCTCGGGTGCTCTG 5′
↓x

5′TCGTCGGCAGCGTCAGATGTGTATAGAGAGACAGATTGCGCAATGNNNNNNNNNWWrGrGrG----(A)$_{30}$CTGTCTCTTATACACATCTCCGAGCCCACGAGAC 3′
3′AGCAGCCGTCGCAGTCTACACATATTCTCTGTCTAACGCGTTACNNNNNNNNNWWCCC ---- NV(T)$_{30}$GACAGAGAATATGTGTAGAGGCTCGGGTGCTCTG 5′

5) Sample index PCR

FWP: 5′AATGATACGGCGACCACCGAGATCTACAC-NNNNNNNN-TCGTCGGCAGCGTC3′
P5 tail Sample index (i5) s5
RVP: 5′CAAGCAGAAGACGGCATACGAGAT-NNNNNNNN-GTCTCGTGGGCTCGG3′
P7 tail Sample index (i7) s7

5′ TCGTCGGCAGCGTCAGATGTGTATAGAGAGACAGATTGCGCAATGNNNNNNNNNWWrGrGrG -(A)$_{30}$CTGTCTCTTATACACATCTCCGAGCCCACGAGAC 3′
← 3′GGCTCGGGTGCTCTG(i5)TAGAGCATACGGCAGAAGACGAAC5′
5′AATGATACGGCGACCACCGAGATCTACAC(i7)TCGTCGGCAGCGTC3′ →
3′ AGCAGCCGTCGCAGTCTACACATATTCTCTGTCTAACGCGTTACNNNNNNNNNWWCCC-NV(T)$_{30}$GACAGAGAATATGTGTAGAGGCTCGGGTGCTCTG 5′

↓
Final 5′ reads for sequencing

5′AATGATACGGCGACCACCGAGATCTACCA(i7)TCGTCGGCAGCGTCAGATGTGTATAAGAGACAGATTGCGCAATGNNNNNNNNNWWrGrGrG-(A)$_{30}$CTGTCTCTTATACACATCTCCGAGCCCACGAGAC(i5)ATCTCGTATGCCGTCTTCTGCTTG 3′
3′TTACTATGCCGCTGGTGGCTCTAGATGTG(i7)AGCAGCCGTCGCAGTCTACACATATTCTCTGTCTAACGCGTTACNNNNNNNNNWW CCC-NV(T)$_{30}$GACAGAGAATATGTGTAGAGGCTCGGGTGCTCTG(i5)TAGAGCATACGGCAGAAGACGAAC 5′

**Supplementary Figure 2.** Selection and isolation of single K562 cell with LCM.

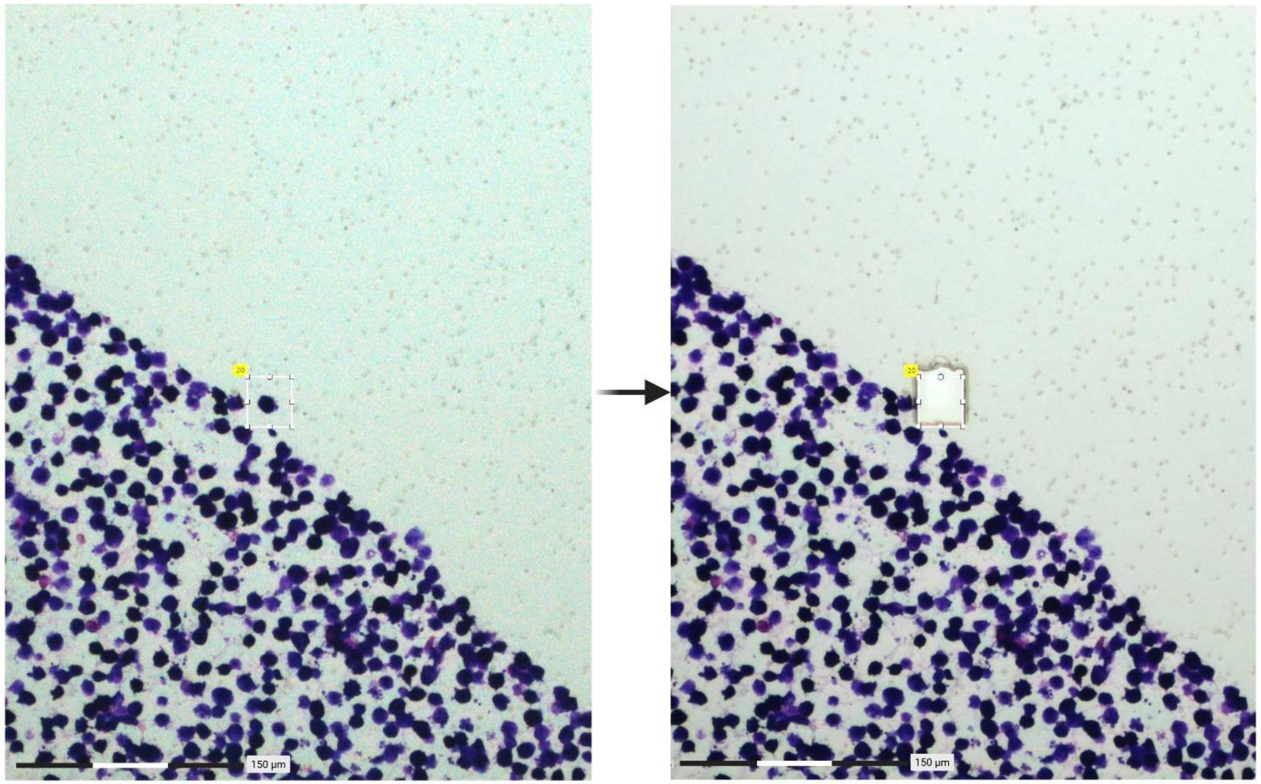


Single K562 cell selection

Isolation of single K562 cell with LCM

**Supplementary Figure 3.** cDNA pre-amplification library Fragment Analyzer profiles of dilution series of LCM-isolated K562 cells. Electropherograms show fragment size distribution and yield following RT and cDNA pre-amplification. LM; Lower Marker, UM; Upper Marker.

| NC 1 | NC 2 | |
|---|---|---|
| 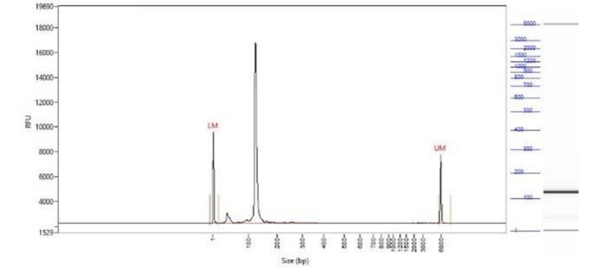 | 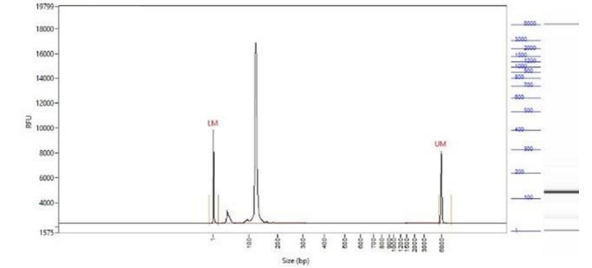 | |
| 1 cell – replicate 1 | 1 cell – replicate 2 | 1 cell – replicate 3 |
| 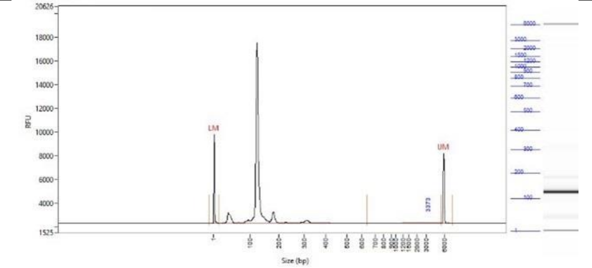 | 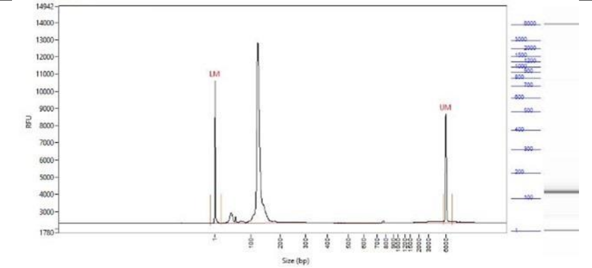 | 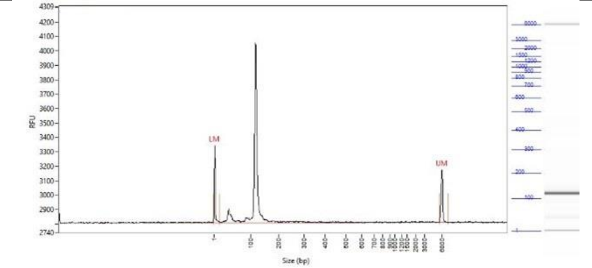 |
| 5 cells – replicate 1 | 5 cells – replicate 2 | 5 cells – replicate 3 |
| 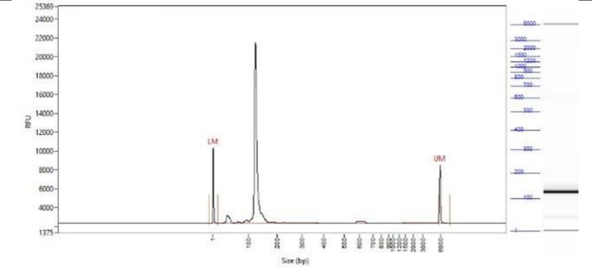 | 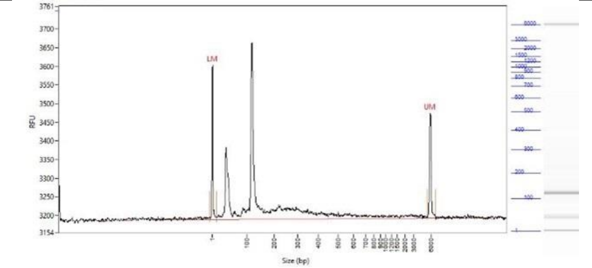 | 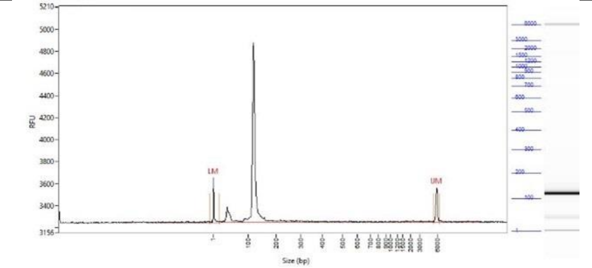 |
| 10 cells – replicate 1 | 10 cells – replicate 2 | 10 cells – replicate 3 |

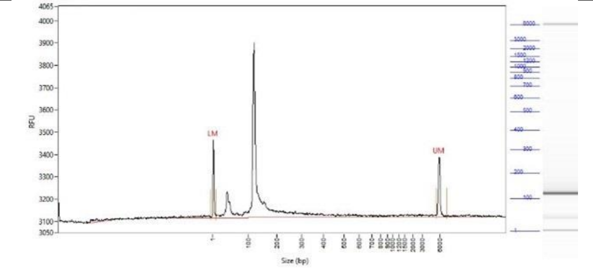

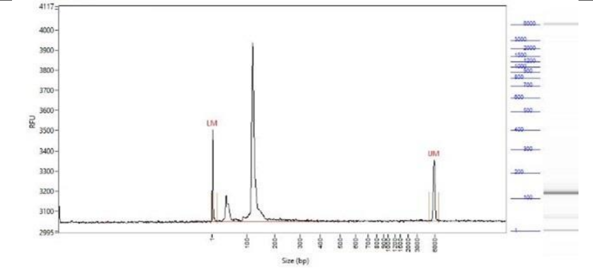

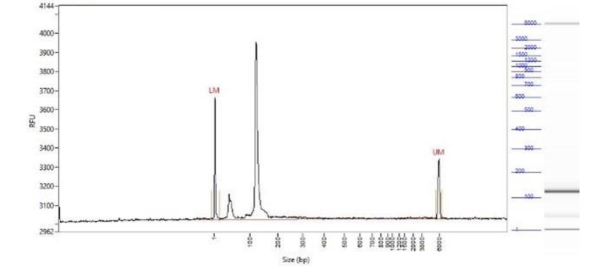


**30 cells – replicate 1** | **30 cells – replicate 2** | **30 cells – replicate 3**

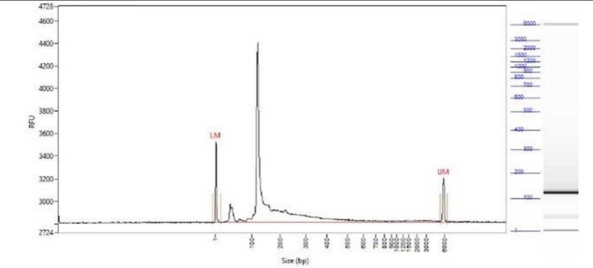

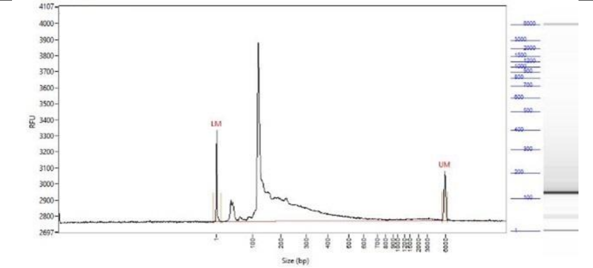

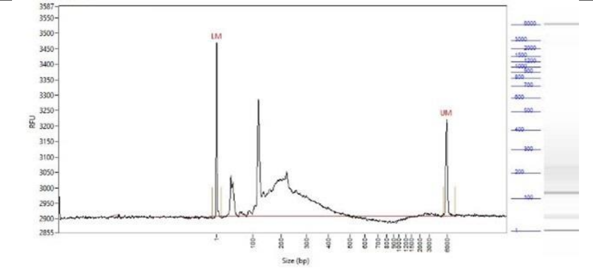


**50 cells – replicate 1** | **50 cells – replicate 2** | **50 cells – replicate 3**

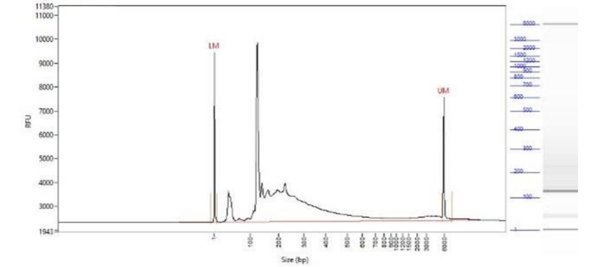

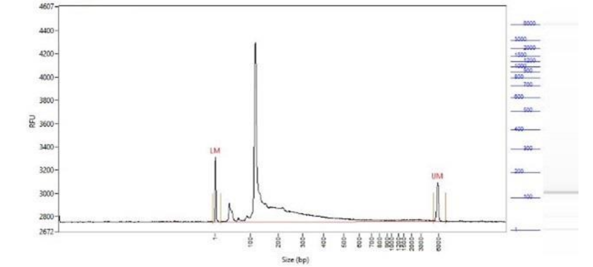

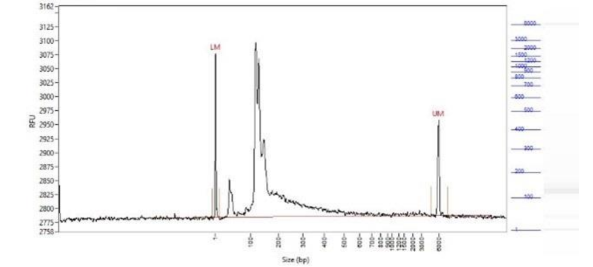


**100 cells – replicate 1** | **100 cells – replicate 2** | **100 cells – replicate 3**

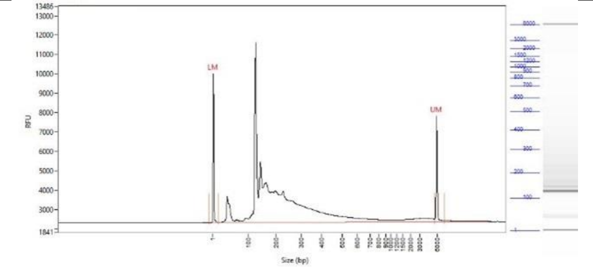

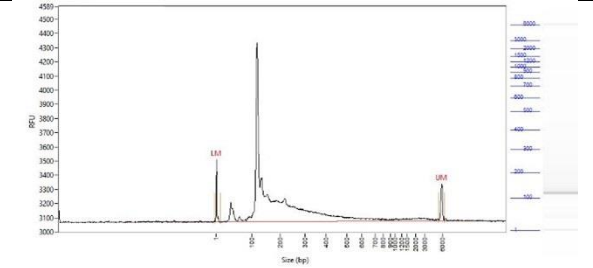

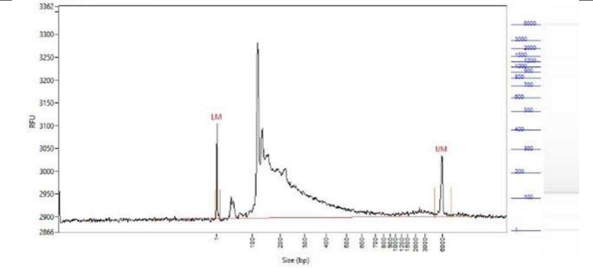

| 300 cells – replicate 1 | 300 cells – replicate 2 | 300 cells – replicate 3 |
| --- | --- | --- |
| 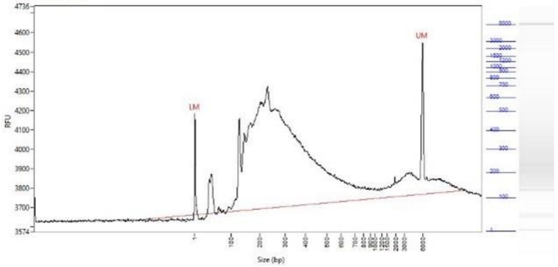 | 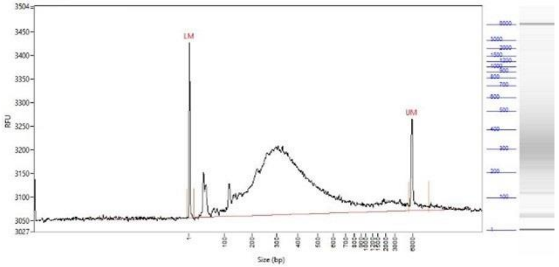 | 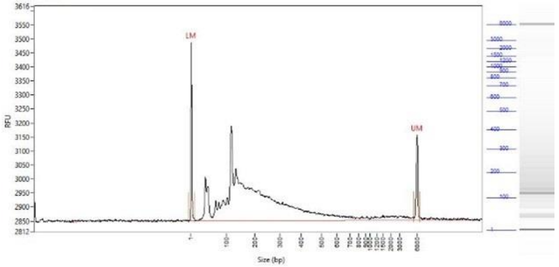 |
| **500 cells – replicate 1** | **500 cells – replicate 2** | **500 cells – replicate 3** |
| 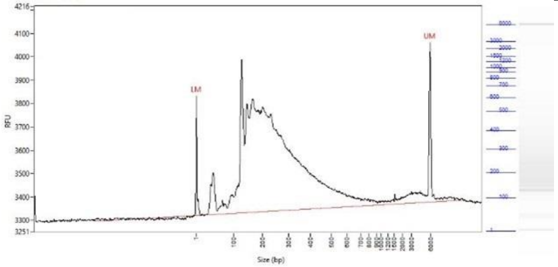 | 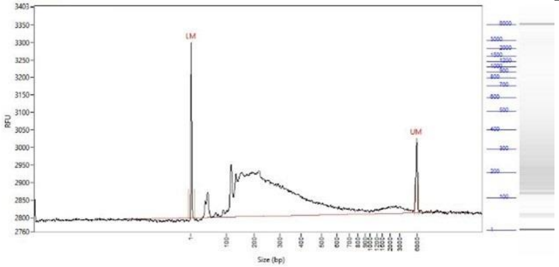 | 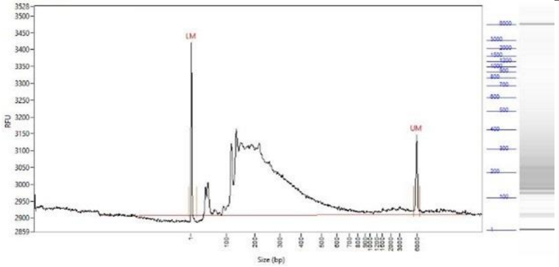 |

**Supplementary Figure 4.** Sequencing saturation curves across the dilution series of LCM-isolated K562 cells. Library complexity was evaluated across the dilution series using saturation curves plotting the detected unique UMI counts against the total number of reads per replicate.

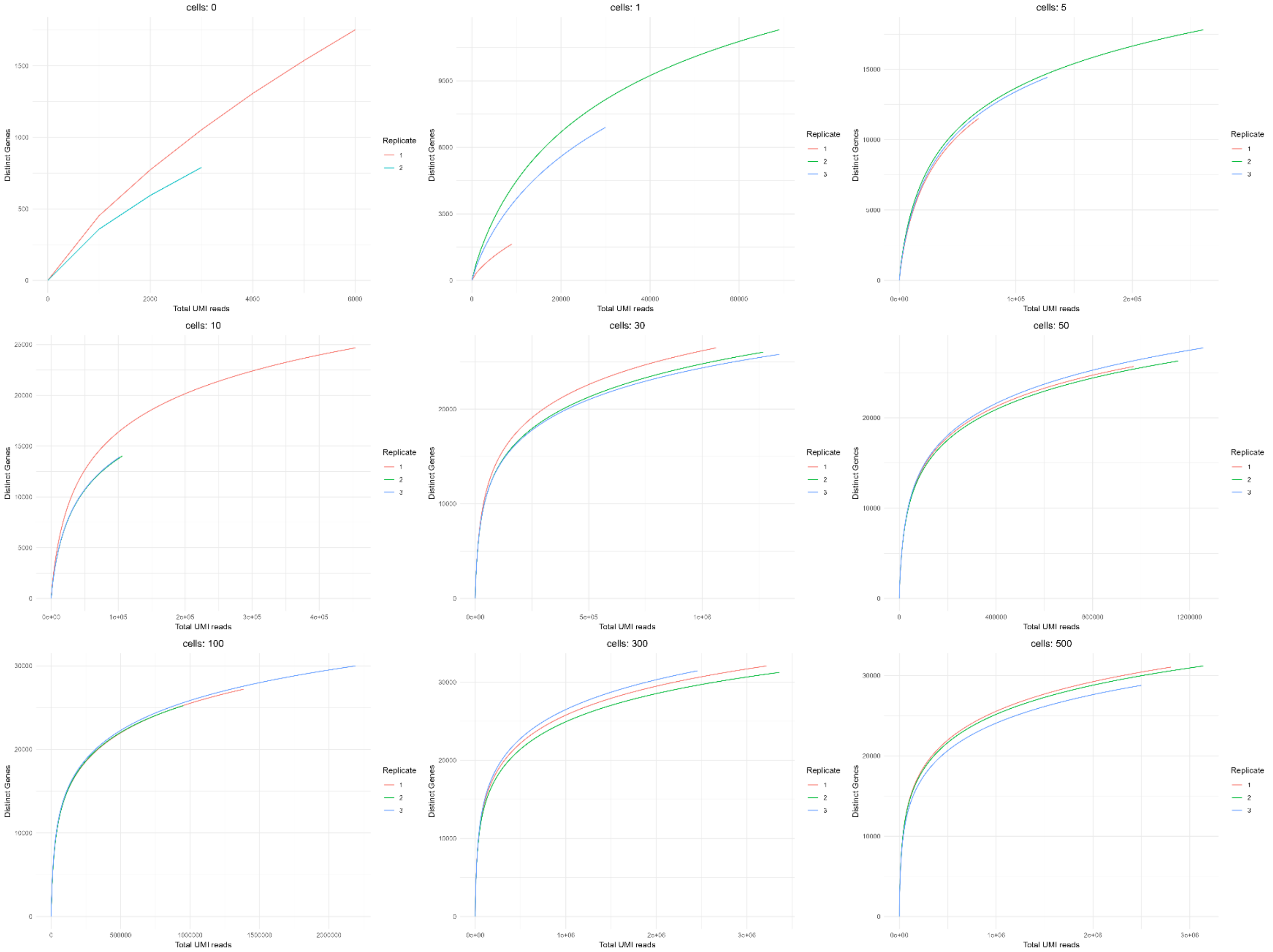

**Supplementary Figure 5.** Selection and isolation of a Sertoli cell with LCM in sample 1 (40x magnification).

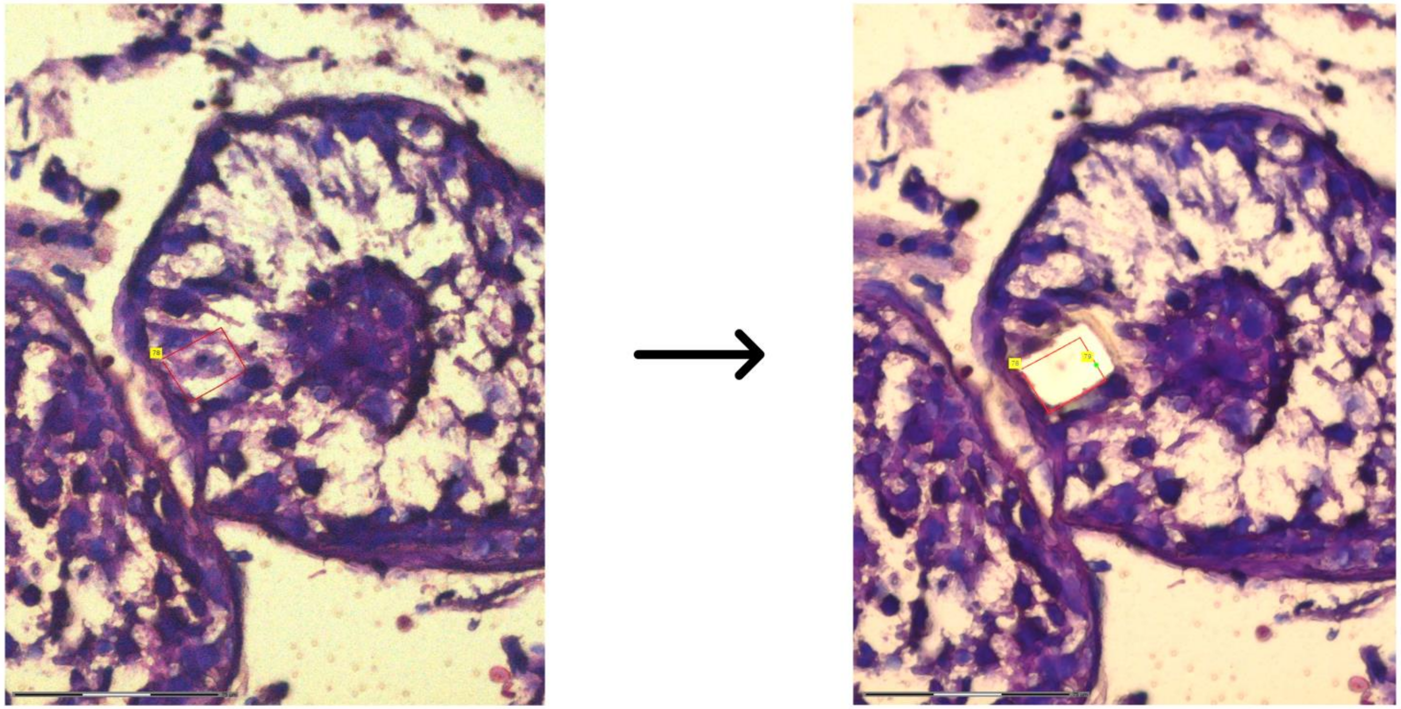

**Supplementary Figure 6.** Proportional average gene biotype distribution per group for dilution series of LCM-isolated Sertoli cells from human control testicular tissue.

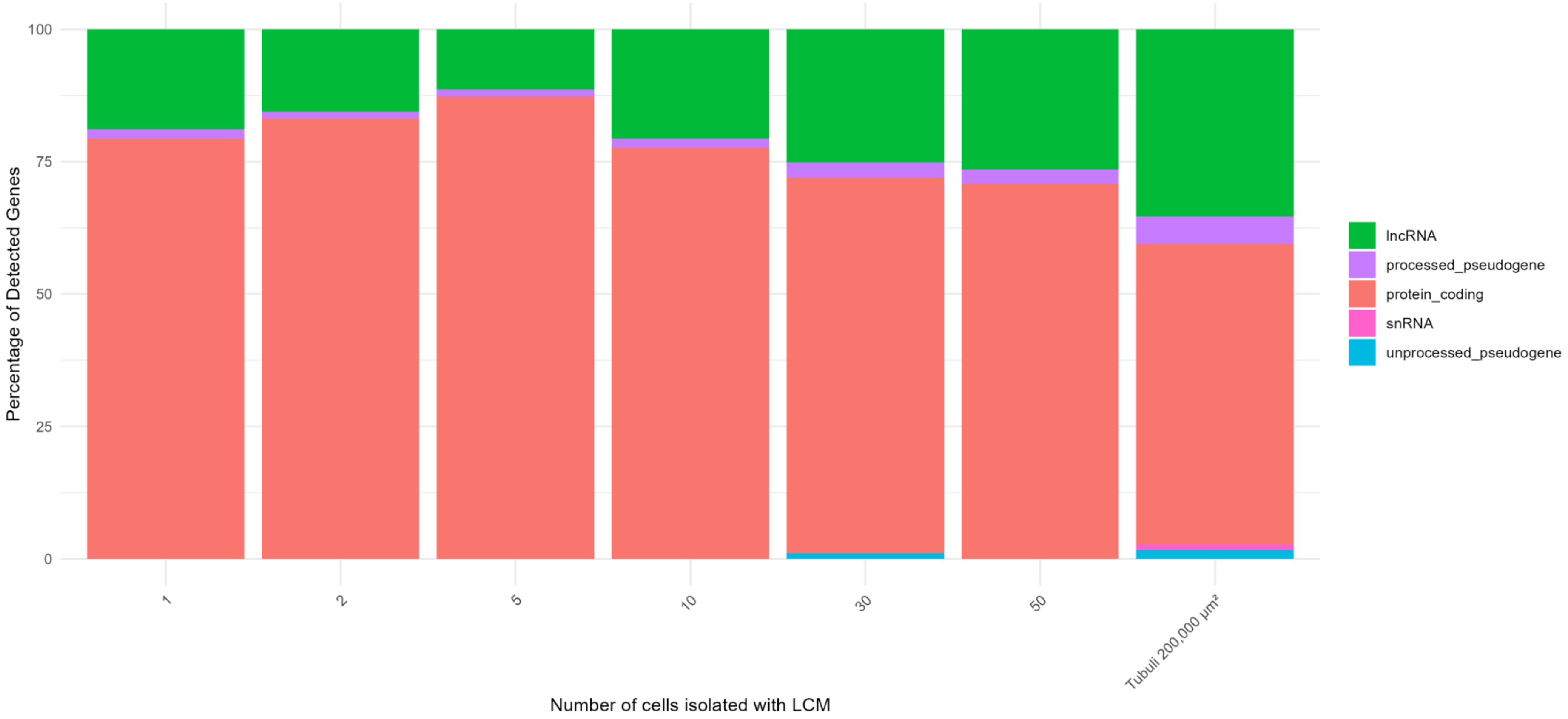

**Supplementary Figure 7.** Sequencing saturation curves across the dilution series of LCM-isolated human Sertoli cells. Library complexity was evaluated across the dilution series using saturation curves plotting the detected unique UMI counts against the total number of reads per replicate.

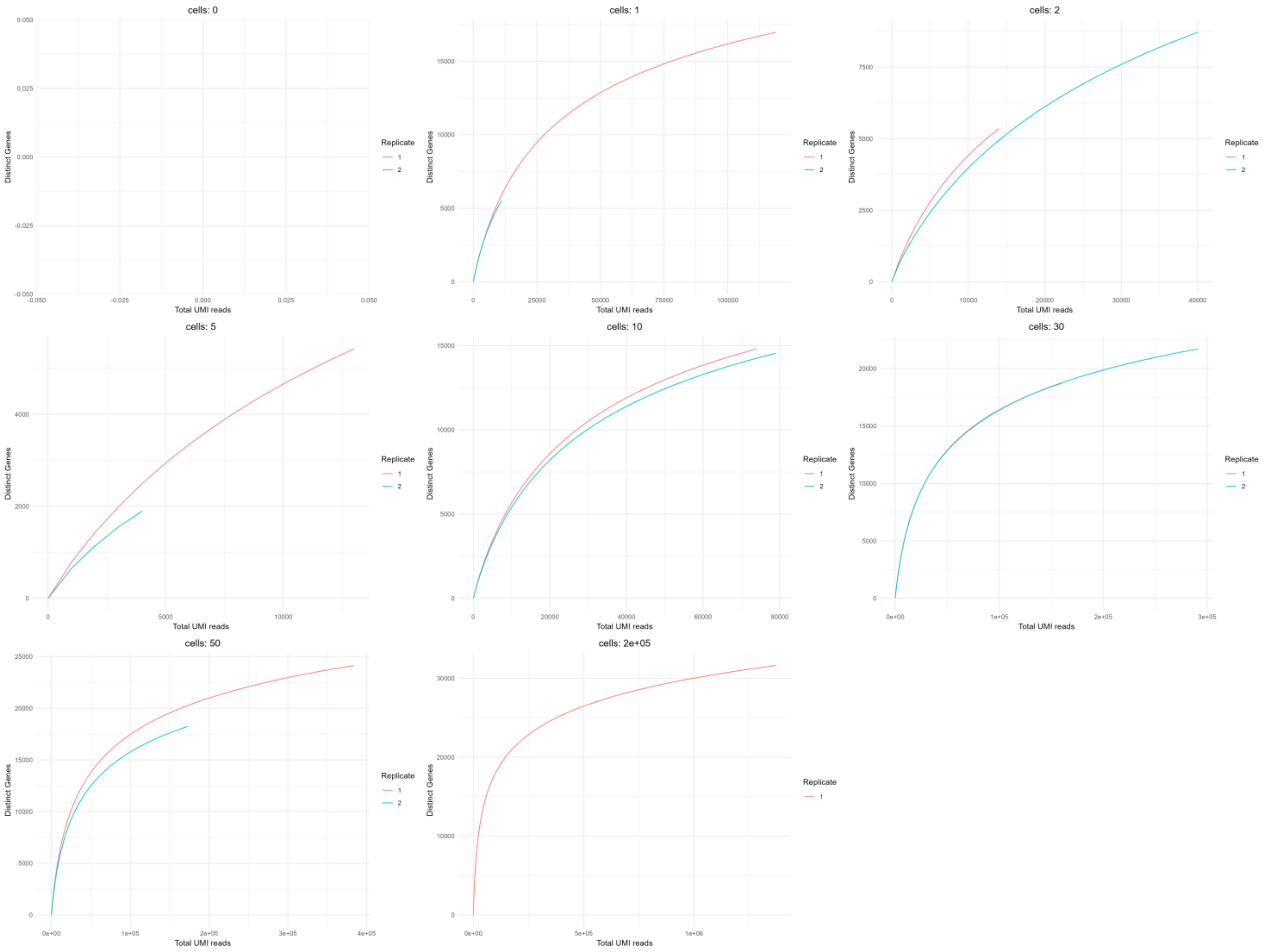

**Supplementary Figure 8.** PCA plot of Sertoli versus Leydig cells. Two-dimensional PCA (based on top 500 most variable genes) demonstrates the clustering of Sertoli cells (triangles) and the clustering of Leydig cells (circles).

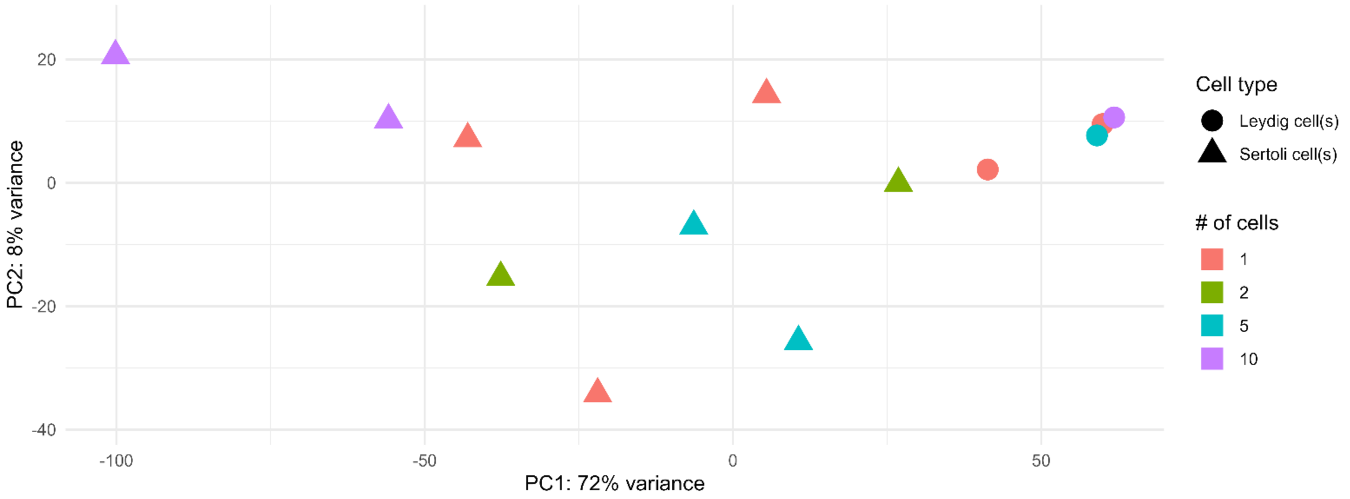

**Supplementary Figure 9.** Volcano plot of DEGs between LCM-isolated human Sertoli versus Leydig cells. The volcano plot displays $log_2FC$ on the x-axis and the $-log10(P_{adj})$ on the y-axis. Single genes are depicted as dots. While more genes were differentially expressed based on $P_{adj} < 0.05$, a total of 328 genes exhibited either upregulation ($log_2FC > 1.5$, shown in red) or downregulation ($log_2FC < -1.5$, shown in blue) in Sertoli cells compared to Leydig cells.

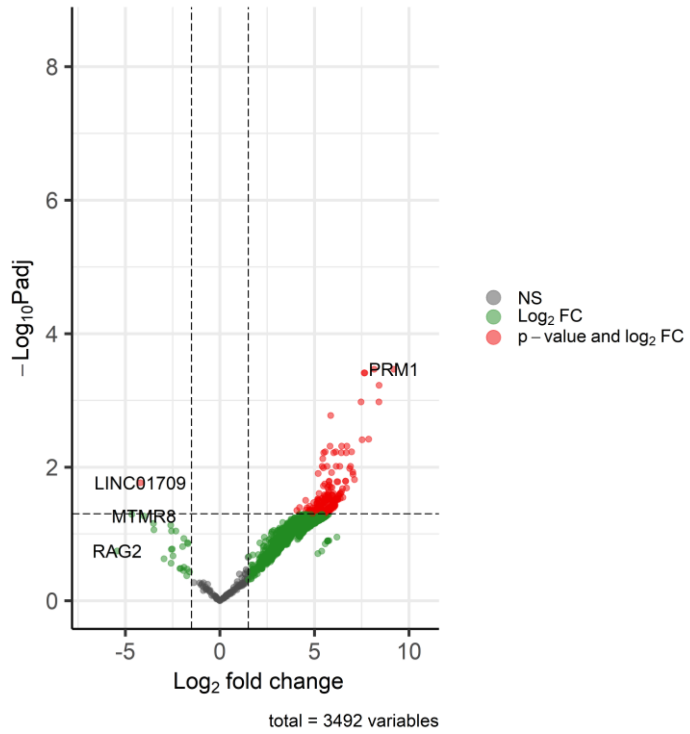

**Supplementary Figure 10.** Average number of detected genes for each sample with seminiferous tubules, shown separately for protein-coding genes and all detected genes. TH: tubular hyalinization.

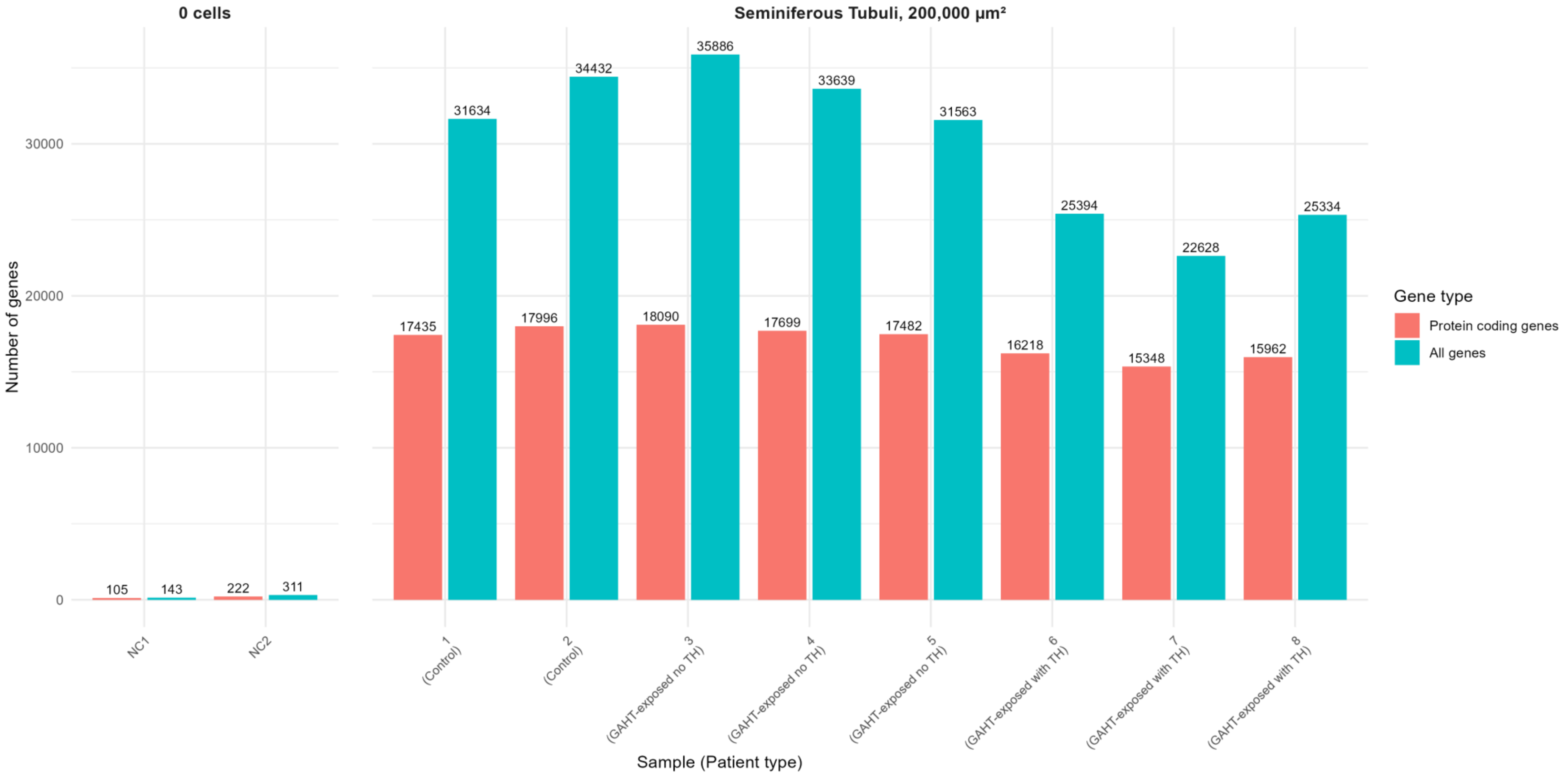

**Supplementary Figure 11.** Volcano plot of DEGs between GAHT-exposed testicular tissue without tubular hyalinization versus controls or GAHT-exposed testicular tissue with tubular hyalinization. The volcano plot displays $log_2FC$ on the x-axis and the -log10($P_{adj}$) on the y-axis. Single genes are depicted as dots. (A) Volcano plot of DEGs between GAHT-exposed testicular tissue without tubular hyalinization versus controls. A total of *1695* genes were significantly downregulated (left side) and *1584* genes significantly upregulated (right side) based on $P_{adj} < 0.05$ and $|log_2FC| > 1.5$. (B) Volcano plot of DEGs between GAHT-exposed testicular tissue with tubular hyalinization versus without tubular hyalinization. A total of *1121* genes were significantly downregulated (left side) and *124* genes significantly upregulated (right side) based on $P_{adj} < 0.05$ and $|log_2FC| > 1.5$.

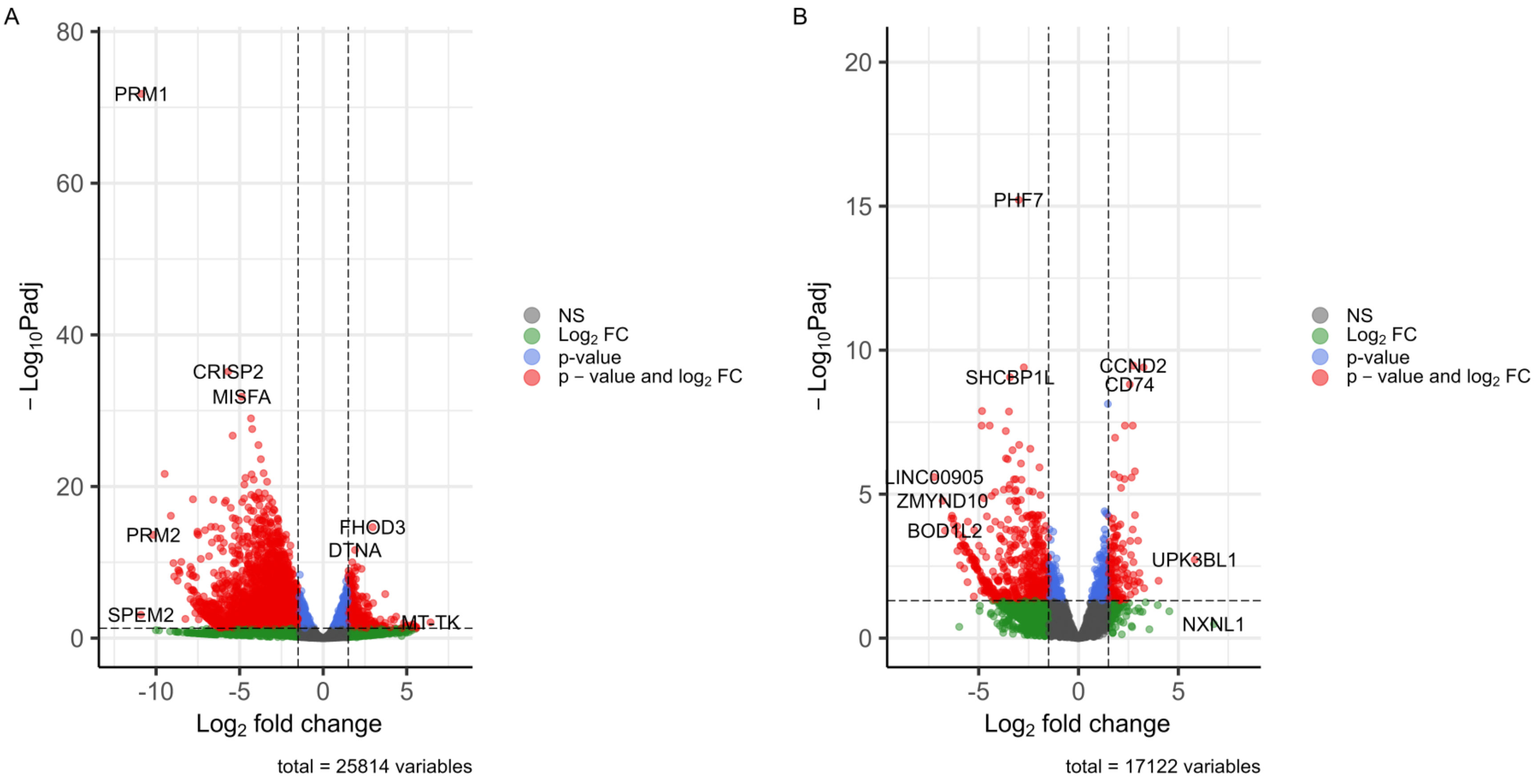

**Supplementary Figure 12**. Gene expressions relative to control samples 1 and 2. Bar plots showing the average log2 expression ratio of gene sets (from Figure 5) compared to control testicular tissue with in A the genes overexpressed in the control tissue and in B the genes overexpressed in the GAHT-exposed testicular tissue.

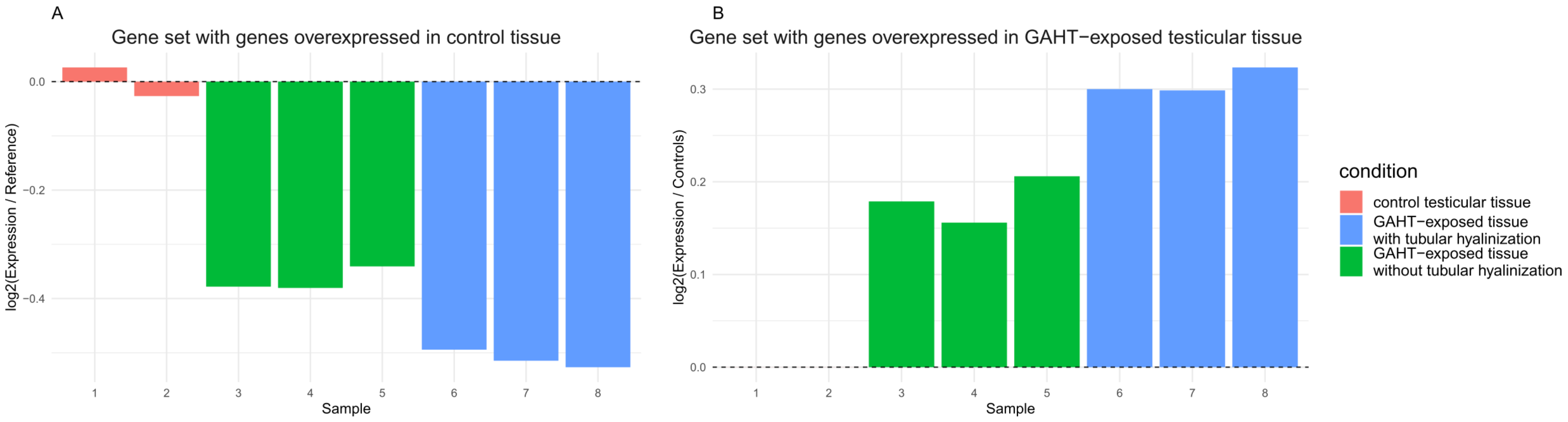

#### Supplementary Tables

**Supplementary Table 1.** Detailed RNA-seq metrics of validation of dilution series of LCM-isolated K562 cells. Sample IDs are indicated as X.Y, where X represents the number of cells isolated with LCM and Y indicates the replicate number.

| | Sample ID | Total | Intergenic reads (%) | Intron reads (%) | Exon reads (%) | Ambiguity reads (%) | Unmapped reads (%) | UMI count | Exon Gene Count (zUMIs) | Protein-coding Gene Count (Biotype) |
|---|---|---|---|---|---|---|---|---|---|---|
| 0 cells | 0.1 | 698,837 | 12.12 | 0.02 | 1.13 | 0.00 | 85.08 | 6,083 | 690 | 1,360 |
| | 0.2 | 430,638 | 14.15 | 0.02 | 1.84 | 0.00 | 82.16 | 3,467 | 209 | 653 |
| 1 cell | 1.1 | 1,016,422 | 6.32 | 0.01 | 0.40 | 0.00 | 92.09 | 9,041 | 422 | 1,252 |
| | 1.2 | 677,047 | 12.15 | 0.05 | 9.72 | 0.00 | 72.57 | 69,636 | 7,496 | 8,981 |
| | 1.3 | 627,159 | 12.11 | 0.04 | 7.73 | 0.00 | 76.16 | 30,402 | 4,335 | 5,724 |
| 5 cells | 5.1 | 887,630 | 11.35 | 0.05 | 9.34 | 0.00 | 74.29 | 68,401 | 7,885 | 9,138 |
| | 5.2 | 2,212,703 | 25.51 | 0.16 | 36.51 | 0.01 | 20.74 | 261,987 | 13,026 | 12,698 |
| | 5.3 | 1,616,143 | 16.06 | 0.08 | 17.31 | 0.00 | 57.81 | 127,756 | 10,259 | 10,974 |
| 10 cells | 10.1 | 5,856,974 | 26.02 | 0.12 | 19.53 | 0.00 | 42.47 | 454,412 | 17,759 | 15,545 |
| | 10.2 | 1,392,421 | 15.53 | 0.06 | 12.15 | 0.00 | 66.16 | 106,520 | 9,765 | 10,728 |
| | 10.3 | 1,591,172 | 12.52 | 0.05 | 10.51 | 0.00 | 71.29 | 102,737 | 9,489 | 10,631 |
| 30 cells | 30.1 | 8,825,296 | 23.27 | 0.15 | 30.45 | 0.01 | 30.85 | 1,061,761 | 19,637 | 15,778 |
| | 30.2 | 6,183,460 | 22.51 | 0.14 | 34.64 | 0.01 | 28.01 | 1,268,205 | 19,657 | 15,649 |
| | 30.3 | 8,810,136 | 22.86 | 0.17 | 43.22 | 0.01 | 16.23 | 1,340,361 | 19,320 | 15,565 |
| 50 cells | 50.1 | 8,204,024 | 25.70 | 0.16 | 35.54 | 0.01 | 22.21 | 970,543 | 18,888 | 15,617 |
| | 50.2 | 9,412,686 | 23.73 | 0.16 | 34.99 | 0.01 | 24.06 | 1,154,720 | 19,546 | 15,684 |
| | 50.3 | 14,143,794 | 21.28 | 0.11 | 20.82 | 0.00 | 46.59 | 1,258,360 | 20,903 | 16,210 |
| 100 cells | 100.1 | 8,244,272 | 24.84 | 0.14 | 35.31 | 0.01 | 25.50 | 1,384,097 | 20,269 | 16,014 |
| | 100.2 | 6,024,094 | 22.85 | 0.13 | 31.17 | 0.01 | 32.14 | 952,005 | 18,589 | 15,462 |
| | 100.3 | 10,720,583 | 23.58 | 0.15 | 31.93 | 0.01 | 29.22 | 2,190,088 | 23,082 | 16,697 |
| 300 cells | 300.1 | 14,772,820 | 23.80 | 0.16 | 36.04 | 0.01 | 23.50 | 3,216,538 | 25,328 | 17,098 |
| | 300.2 | 12,086,787 | 23.64 | 0.15 | 38.95 | 0.01 | 21.61 | 3,358,456 | 24,553 | 16,840 |
| | 300.3 | 9,901,552 | 26.57 | 0.12 | 33.19 | 0.01 | 27.82 | 2,451,933 | 24,495 | 17,007 |
| 500 cells | 500.1 | 9,231,132 | 25.78 | 0.14 | 33.51 | 0.01 | 25.51 | 2,804,343 | 24,245 | 16,828 |
| | 500.2 | 10,453,342 | 24.72 | 0.14 | 34.52 | 0.01 | 25.81 | 3,140,756 | 24,478 | 16,912 |
| | 500.3 | 6,887,198 | 23.37 | 0.16 | 39.62 | 0.01 | 20.47 | 2,499,420 | 22,200 | 16,322 |

| Successful samples | | | | | | | | | |
|---|---|---|---|---|---|---|---|---|---|
| MEAN | 6,902,714 | 21.29 | 0.12 | 27.68 | 0.01 | 38.31 | 1,316,236 | 17,618 | 14,265 |
| MIN | 627,159 | 11.35 | 0.04 | 7.73 | 0.00 | 16.23 | 30,402 | 4,335 | 5,724 |
| MAX | 14,772,820 | 26.57 | 0.17 | 43.22 | 0.01 | 76.16 | 3,358,456 | 25,328 | 17,098 |

| Successful single-cell samples | | | | | | | | | |
|---|---|---|---|---|---|---|---|---|---|
| MEAN | 652,103 | 12.13 | 0.05 | 8.73 | 0.00 | 74.36 | 50,019 | 5,916 | 7,353 |
| MIN | 627,159 | 12.11 | 0.04 | 7.73 | 0.00 | 72.57 | 30,402 | 4,335 | 5,724 |
| MAX | 677,047 | 12.15 | 0.05 | 9.72 | 0.00 | 76.16 | 69,636 | 7,496 | 8,981 |

| 30-500 cells | | | | | | | | | |
|---|---|---|---|---|---|---|---|---|---|
| MEAN | 9,593,412 | 24 | 0.15 | 34 | 0 | 27 | 1,936,772 | 21,679 | 16,246 |
| MIN | 6,024,094 | 21 | 0.11 | 21 | 0 | 16 | 952,005 | 18,589 | 15,462 |
| MAX | 14,772,820 | 27 | 0.17 | 43 | 0 | 47 | 3,358,456 | 25,328 | 17,098 |

**Supplementary Table 2.** Specifications of dilution series of LCM-isolated Sertoli cells and corresponding cDNA libraries. Values are mean ± SD. NA, not applicable.

| **Number of LCM-isolated cells** | **Replicates** | **Cells collected** | **Total Area (µm²)** | **cDNA yield (ng)** |
|---|---|---|---|---|
| 0 cells | 2 | NA | NA | 5.6 ± 1.3 |
| 1 cell | 3 | 1 ± 0 | 428 ± 48 | 10.4 ± 6.1 |
| 2 cells | 2 | 2 ± 0 | 599 ± 163 | 10.8 ± 0.1 |
| 5 cells | 2 | 5 ± 0 | 1,266 ± 1,106 | 9.3 ± 2.2 |
| 10 cells | 2 | 10 ± 0 | 1,747 ± 425 | 6.6 ± 1.8 |
| 30 cells | 2 | 30 ± 0 | 2,249 ± 1,135 | 7.8 ± 3.6 |
| 50 cells | 2 | 50 ± 0 | 3,262 ± 297 | 12.3 ± 2.8 |

**Supplementary Table 3.** Detailed RNA-seq metrics of validation of dilution series of LCM-isolated Sertoli cells, Leydig cells and seminiferous tubules samples. Sample IDs are indicated as X.Y, where X represents the number of cells isolated with LCM and Y indicates the replicate number.

**Dilution Series of LCM-isolated Sertoli Cells**

| | Sample ID | Total | Intergenic reads (%) | Intron reads (%) | Exon reads (%) | Ambiguity reads (%) | Unmapped reads (%) | UMI count | Exon Gene Count zUMIs | Gene Count Biotype Protein-Coding Genes only |
|---|---|---|---|---|---|---|---|---|---|---|
| 0 cells | 0.1 | 149,478 | 2.33 | 0.16 | 0.10 | 0.01 | 97.41 | 322 | 30 | 105 |
| | 0.2 | 513,944 | 2.27 | 0.16 | 0.10 | 0.00 | 97.46 | 735 | 63 | 222 |
| 1 cell | 1.1 | 7,504,627 | 21.37 | 9.76 | 37.34 | 0.70 | 30.83 | 119,863 | 13,276 | 12,668 |
| | 1.2 | 4,345,888 | 10.84 | 3.01 | 3.18 | 0.09 | 82.88 | 11,155 | 2,211 | 4,111 |
| | 1.3 | 1,026,381 | 13.48 | 3.97 | 24.47 | 0.51 | 57.57 | 7,383 | 2,267 | 2,692 |
| 2 cells | 2.1 | 1,909,762 | 10.29 | 5.06 | 18.86 | 0.36 | 65.43 | 14,133 | 3,460 | 4,532 |
| | 2.2 | 4,566,427 | 8.09 | 3.71 | 4.03 | 0.10 | 84.08 | 40,880 | 4,686 | 6,901 |
| 5 cells | 5.1 | 1,689,846 | 13.12 | 5.48 | 26.18 | 0.58 | 54.65 | 13,369 | 3,930 | 4,680 |
| | 5.2 | 583,312 | 9.73 | 3.97 | 15.92 | 0.18 | 70.20 | 4,113 | 1,322 | 1,652 |
| 10 cells | 10.1 | 3,937,258 | 25.93 | 11.03 | 49.61 | 1.11 | 12.31 | 74,878 | 11,803 | 11,162 |
| | 10.2 | 3,489,166 | 13.42 | 6.97 | 29.13 | 0.60 | 49.88 | 79,397 | 11,269 | 11,165 |
| 30 cells | 30.1 | 5,291,191 | 18.30 | 12.16 | 27.93 | 0.56 | 41.07 | 160,687 | 14,343 | 13,427 |
| | 30.2 | 3,853,329 | 21.36 | 12.60 | 27.53 | 0.72 | 37.79 | 291,876 | 17,362 | 14,474 |
| 50 cells | 50.1 | 9,108,233 | 22.83 | 12.56 | 42.96 | 1.02 | 20.63 | 384,039 | 19,988 | 15,547 |
| | 50.2 | 6,106,991 | 23.69 | 15.61 | 50.54 | 1.05 | 9.11 | 173,426 | 14,632 | 13,306 |

| Successful samples | | | | | | | | | |
|---|---|---|---|---|---|---|---|---|---|
| **MEAN** | 4,108,647 | 16.34 | 8.14 | 27.51 | 0.58 | 47.42 | 105,785 | 9,273 | 8,947 |
| **MIN** | 583,312 | 8.09 | 3.01 | 3.18 | 0.09 | 9.11 | 4,113 | 1,322 | 1,652 |
| **MAX** | 9,108,233 | 25.93 | 15.61 | 50.54 | 1.11 | 84.08 | 384,039 | 19,988 | 15,547 |

| Successful single-cell samples | | | | | | | | | |
|---|---|---|---|---|---|---|---|---|---|
| **MEAN** | 4,292,299 | 15.23 | 5.58 | 21.66 | 0.43 | 57.09 | 46,134 | 5,918 | 6,490 |
| **MIN** | 1,026,381 | 10.84 | 3.01 | 3.18 | 0.09 | 30.83 | 7,383 | 2,211 | 2,692 |
| **MAX** | 7,504,627 | 21.37 | 9.76 | 37.34 | 0.70 | 82.88 | 119,863 | 13,276 | 12,668 |

| 30-50 cells | | | | | | | | | |
|---|---|---|---|---|---|---|---|---|---|
| **MEAN** | 6,089,936 | 21.54 | 13.23 | 37.24 | 0.84 | 27.15 | 252,507 | 16,581 | 14,189 |
| **MIN** | 3,853,329 | 18.30 | 12.16 | 27.53 | 0.56 | 9.11 | 160,687 | 14,343 | 13,306 |
| **MAX** | 9,108,233 | 23.69 | 15.61 | 50.54 | 1.05 | 41.07 | 384,039 | 19,988 | 15,547 |

**Leydig Cells**

| Sample ID | Total | Intergenic reads (%) | Intron reads (%) | Exon reads (%) | Ambiguity reads (%) | Unmapped reads (%) | UMI count | Exon Gene Count zUMIs | Gene Count Biotype Protein-Coding Genes only |
|---|---|---|---|---|---|---|---|---|---|
| 1.1 | 254,929 | 2.13 | 0.53 | 0.25 | 0.02 | 97.07 | 903 | 113 | 232 |
| 1.2 | 1,861,374 | 14.78 | 3.07 | 8.33 | 0.12 | 73.70 | 17,040 | 3,476 | 4,714 |
| 5.1 | 259,615 | 4.66 | 0.44 | 0.32 | 0.00 | 94.57 | 1,402 | 118 | 281 |
| 10.1 | 198,106 | 4.35 | 0.57 | 0.10 | 0.00 | 94.97 | 1,011 | 59 | 175 |

| | | | | | | | | | |
|---|---|---|---|---|---|---|---|---|---|
| **MEAN** | 643,506 | 6.48 | 1.15 | 2.25 | 0.04 | 90.08 | 5,089 | 942 | 1,351 |
| **MAX** | 1,861,374 | 14.78 | 3.07 | 8.33 | 0.12 | 97.07 | 17,040 | 3,476 | 4,714 |
| **MIN** | 198,106 | 2.13 | 0.44 | 0.10 | 0.02 | 73.7 | 903 | 59 | 175 |

**Seminiferous Tubules Samples**

| | Sample ID | Total | Intergenic reads (%) | Intron reads (%) | Exon reads (%) | Ambiguity reads (%) | Unmapped reads (%) | UMI count | Exon Gene Count zUMIs | Gene Count Biotype Protein-Coding Genes only |
|---|---|---|---|---|---|---|---|---|---|---|
| Controls - no tubular hyalinization, 200,000 µm² | 1 | 7,293,748 | 25.39 | 12.78 | 41.36 | 1.15 | 19.32 | 1,371,567 | 27,852 | 17,435 |
| | 2 | 7,355,356 | 27.77 | 12.71 | 38.24 | 1.01 | 20.27 | 2,206,306 | 30,445 | 17,996 |
| GAHT-exposed - no tubular hyalinization, 200,000 µm² | 3 | 10,570,421 | 35.55 | 13.84 | 32.03 | 0.57 | 18.02 | 3,366,711 | 30,905 | 18,090 |
| | 4 | 8,937,026 | 39.02 | 13.48 | 12.84 | 0.33 | 34.33 | 1,583,472 | 27,656 | 17,699 |
| | 5 | 11,632,685 | 20.25 | 12.48 | 34.52 | 0.61 | 32.15 | 1,900,034 | 27,302 | 17,482 |
| GAHT-exposed – with tubular hyalinization, 200,000 µm² | 6 | 8,889,581 | 25.16 | 19.77 | 34.33 | 0.80 | 19.94 | 657,856 | 20,339 | 16,218 |
| | 7 | 9,375,724 | 21.09 | 14.45 | 53.28 | 0.93 | 10.25 | 524,799 | 18,400 | 15,348 |
| | 8 | 10,522,404 | 18.29 | 15.42 | 38.63 | 0.78 | 26.88 | 853,651 | 20,187 | 15,962 |

| | | | | | | | | | |
|---|---|---|---|---|---|---|---|---|---|
| **MEAN** | 9,364,114 | 26.33 | 14.54 | 37.08 | 0.80 | 21.26 | 1,488,800 | 25,144 | 17,007 |
| **MIN** | 7,293,748 | 18.29 | 12.48 | 12.84 | 0.33 | 10.16 | 524,799 | 18,400 | 15,348 |
| **MAX** | 11,632,685 | 39.02 | 19.77 | 53.28 | 1.15 | 34.33 | 3,366,711 | 30,905 | 18,090 |
